# Viral transduction and the dynamics of bacterial adaptation

**DOI:** 10.1101/2021.06.14.448024

**Authors:** Philippe Cherabier, Sylvie Méléard, Régis Ferrière

**Affiliations:** Institut de Biologie de l’École Normale Supérieure (IBENS), Université Paris Sciences et Lettres, CNRS, INSERM, 75005 Paris, France; CMAP, CNRS, École polytechnique, Institut polytechnique de Paris F-91128 Palaiseau, France et Institut Universitaire de France; Department of Ecology & Evolutionary Biology, University of Arizona, Tucson, AZ 85721, USA; International Research Laboratory for Interdisciplinary Global Environmental Studies (iGLOBES), CNRS, ENS-PSL University, University of Arizona, Tucson, AZ 85721, USA

## Abstract

Transduction - horizontal gene transfer (HGT) by viruses - is an important macroevolutionary force in prokaryotes, contributing to functional innovation and lineage diversification. In contrast, the role that transduction plays in bacterial microevolutionary adaptation is poorly known. By facilitating the transfer of beneficial alleles between host cells, transduction may accelerate adaptation. But transduction also carries the risk of transferring deleterious alleles, which, in addition to the demographic cost of viral infection, may hinder adaptation. Here we resolve the conflicting effects of transduction on bacterial adaptation in a simple eco-evolutionary model for large populations characterized by a quantitative (resource-use) trait with a single evolutionary optimum. Our model focuses on generalized transduction by virulent phages. Away from the optimum, the effect of transferring beneficial alleles dominates and transduction tends to accelerate adaptation. Close to the optimum, transduction generates a large amount of stochasticity in the population adaptive trajectory, thus hindering adaptation. Under disruptive selection, transduction may either limit (as sexual recombination would) or promote phenotypic diversification, or drive ‘transient optimization’ whereby phenotypic subpopulations recurrently visit the optimum. Our modeling framework paves the way to study complex adaptive feedbacks between bacterial hosts and phages generated by the combination of deterministic and stochastic effects of transduction.

## Introduction

Bacteria can transfer genetic material ‘vertically’ to daughter cells as well as ‘horizontally’ between cells. The process of horizontal gene transfer allows bacterial species to successfully conquer and adapt to new ecological niches (De la Cruz and Davies 2000, Ochman et al. 2000, Lawrence 2002) by driving the rapid spread of evolutionary innovation (Hall et al. 2017, Hao and Golding 2006). Well-studied mechanisms of horizontal gene transfer (HGT) between bacterial cells are transformation, conjugation, and transduction. Of the three, transduction by bacterial viruses, or phages, is generally regarded as the most important (Chen et al. 2018). Transduction is the process whereby host DNA is packaged into phage particles which then deliver the bacterial DNA as they infect other cells. Through transduction, viruses can influence the genotypic composition of host populations, hence potentially their hosts’ evolution. For example, phages are notorious for their role in the transfer of pathogenicity islands and antibiotic resistance genes.

While transduction may contribute substantially to the evolution of major ecological innovations in prokaryotes, very little is known about the role that transduction plays in the adaptation of bacterial populations to small or gradual, quantitative, rather than large, qualitative, changes in their environment (Raz and Tannenbaum 2010, Touchon et al. 2017). The conventional view is that, as a mechanism that facilitates the spread of genetic elements among host cells, transduction may accelerate bacterial adaptation to a changing environment (Andam et al. 2015). But if bacteria can acquire advantageous mutations through transduction, they may also receive deleterious ones (Redfield 1988, Billiard et al. 2016). In addition, if a high rate of transduction comes at the cost of a high rate of viral infection, the adaptive benefit of spreading beneficial mutations may be further eroded by the overall mortality cost of infection. Our goal here is to resolve these conflicting effects of transduction on bacterial adaptation by developing and analysing a simple mathematical model.

Different mechanisms of transduction have been described, depending on the phage life cycle and the region of host DNA that can be transferred (Fineran et al. 2009; Davidson 2018). Specialized transduction and lateral transduction involve temperate phages, i.e. phages that can enter either the lytic or lysogenic life cycles. Specialized transduction occurs when a prophage (phage in lysogeny) excises incorrectly from the host bacterial genome and ends up packaging some of the flanking regions of host DNA. The recently described lateral transduction (Chen et al. 2018) also involves temperate phages, whose late excision and *in situ* replication leads to highly efficient packaging of host DNA over several hundred kilobases downstream of the integration site. Finally, generalized transduction can involve phages that are temperate as well as virulent, i.e. that can only replicate via the lytic cycle. In generalized transduction, phages randomly package host DNA instead of their own and thus can transfer any fragment of the bacterial genome. Phages packed with host DNA, or transducing particles, are released during lysis, together with functional virions. Transducing particles have the ability to adsorb to bacterial surface receptors and inject their DNA. However, as they lack viral genes, transducing particles cannot trigger lysis. Instead, the injected bacterial DNA might be integrated into the recipient cell’s genome by recombination. In this study, we focus on generalized transduction by virulent phages, which allows us to keep the mathematical framework simple and tractable.

In our model the phenotype of asexually reproducing bacterial cells is characterized by a quantitative trait which measures the cell’s ability to acquire a single resource. We use a simple genotype-phenotype map based on an infinite-site, infinite allele model in which mutations occur upon cell division and cause small, random phenotypic change. The mutated DNA of a mutant cell may be transferred to recipient cells and integrated in their genome via generalized transduction due to a non-evolving virulent phage. To model selection on bacterial trait variation, we take an eco-evolutionary approach in which the intensity and direction of selection is not given *a priori* (Dieckmann and Régis Ferrière 2004). Rather, selection results from the ecological process of cells competing for resources, with competition being more intense among cells that are phenotypically more similar (Doebeli and Dieckmann 2000).

Starting with an individual-level model of bacteria competing with each other and susceptible to infection by a virulent phage, we extend the model to track the effect of generalized transduction on the cells’ genotype and phenotype. Assuming the existence of a single trait value that maximizes individual resource acquisition (i.e., a single evolutionary optimum), the adaptation process unfolds as the bacterial population moves in its one-dimensional phenotypic space. By assuming that the process is mutation-limited, we derive the general equation driving the adaptive trait dynamics. We combine the mathematical analysis of this equation with numerical simulations of the individual-based process to resolve the effect of generalized transduction on (i) the speed of adaptation away from the evolutionary optimum, (ii) the dynamics of adaptation near the optimum, (iii) the maintenance of phenotypic diversity that may result from disruptive selection around the optimum.

## Model

### Individual-level model of infection and generalized transduction

Our model of generalized transduction is summarized in Figure 1. During the reproduction and encapsulation of the viral genome, DNA ‘mispackaging’ may occur resulting in viral particles that contain part of the host genome instead of viral DNA (Touchon et al. 2017). These viral particles, that we call *Gene Transducing Particles*, or GTP, may no longer cause the lysis of the cells they infect. Instead, they pass genetic material from the previous host on to the receiving cell, where it may be integrated in the genome by recombination. As a consequence, the phenotype of the receiving cell, and its offspring lineage, may be altered (Figure 1a).

**Figure 1.**
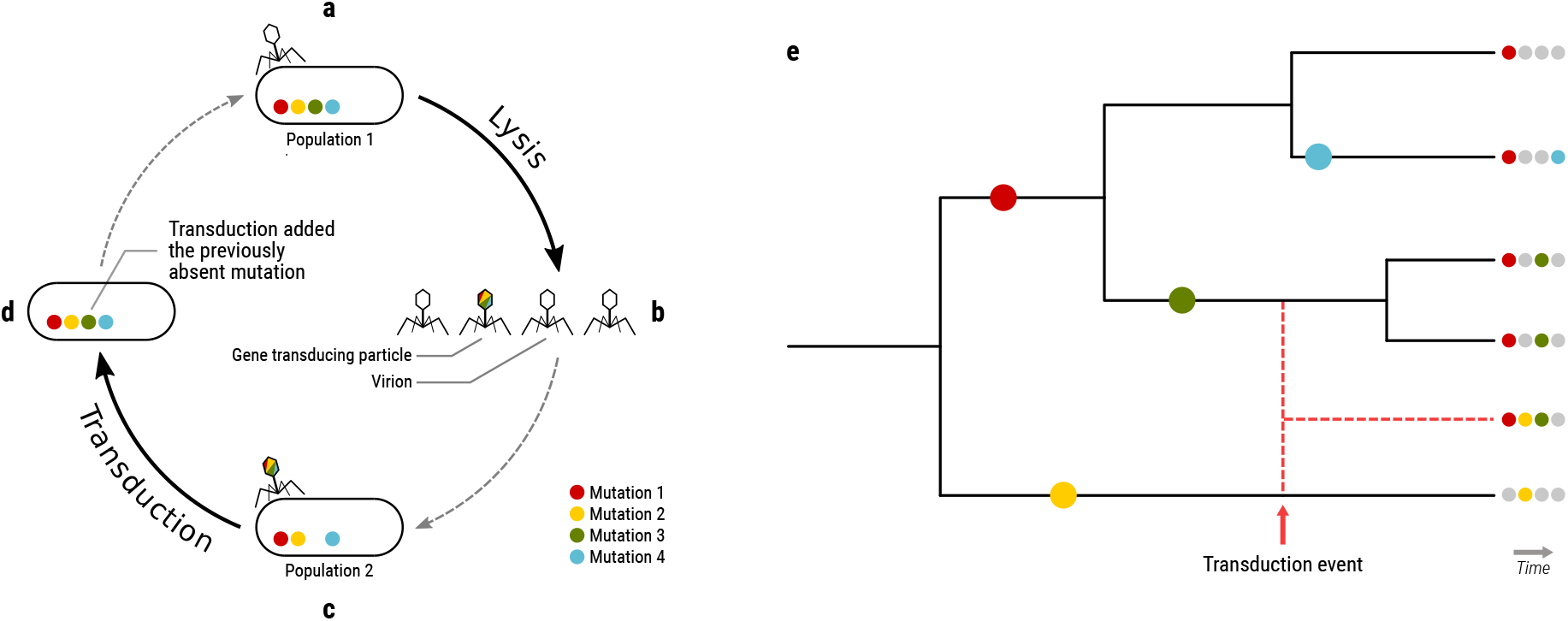
Transduction moves mutant alleles among individuals and create new genotypes. Infection by a lytic virus (a) causes a viral burst. Some of the viral particles released from the burst are genetic transducing particles (b) that may transfer a mutated allele from the previous host to a new host (c). With a given probability of transduction, the mutated allele is integrated by recombination into the new host’s genome (d). Mutations that occur in different lineages are vertically inherited (e). When a transduction event occurs, a new lineage is created (e, dashed red lines).

To describe the genotypic and phenotypic effects of transduction, we focus on a quantitative character, or trait, *a*, and introduce a simple model of the genotype-phenotype map. We make the following assumptions:

1. The trait is under the control of many loci of small, additive effects.
2. We use the principle of an infinite site model, so that no two mutations will impact the same locus.
3. Alleles at the different loci can be moved by transduction. Transduced DNA is integrated by non-homologous end joining (Popa, Hazkani-Covo, et al. 2011), i.e. the new DNA fragment is *added* to the host genome, and does not replace the resident allele.
4. We assume no dosage effect, meaning that the number of occurrences of an allele does not influence the level of expression of the gene.
5. Mispackaging of host DNA in a new viral particle (GTP), the delivery of the previous host’s DNA by the GTP, and the integration of transferred genetic material into the newly infected host’s genome, are stochastic events. Each event occurs with a certain probability.
6. The integration step of the transduction process is more likely if the phenotypes are more similar. The reason is that the underlying genotypic difference between more similar phenotypes is likely smaller, which may facilitate recombination.

Per assumptions 3 and 4, when a cell receives a copy of a mutation it already carries, two copies of the same mutated sequence will occur in the genotype, but assumption 4 implies that the phenotype will not be altered by the second copy of the mutation. Only the presence or absence of an allele influences the cell’s phenotype. The genotype-phenotype map can thus be described as follows. By *n* we denote the number of mutations that have occurred in a population at time *t*. Since each mutation occurs on a different site (assumption 1), we can define 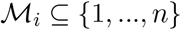 the subset of mutations that individual *i* carries. The phenotypic effect of each mutation (indexed by *m*, varying from 1 to *n*) is a small perturbation *ϵ_m_*. As a consequence, the trait *a_i_* for individual *i* is

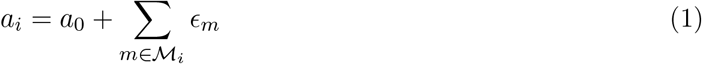

where *a*_0_ is the initial value of the trait in an isogenic population.

Now if *i* and *j* designate two individuals respectively carrying the mutation sets 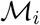 and 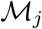, we consider the case where individual *i* encounters a GTP originating from individual *j*, thus carrying 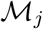. By injecting DNA from individual *j* into the individual *i*, the GTP transfers individual *j*’s specific mutations to the recipient cell’s genotype without altering the mutations already carried by *i*. Thus, i’s mutation set will change from 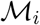 to 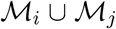, and its phenotype will be altered accordingly. The population coalescent (Figure 1b) shows that when a transduction event takes place between two distinct lineages, a new lineage results from the recombination of the two original genetic material. In that sense, transduction can be seen as driving ‘sparse sexual reproduction events’ (Baltrus et al. 2008). Note that modeling the phenotypic effect of transduction cannot be based solely on individuals’ phenotypes; the explicit tracking of genotypes is required. Indeed, if 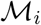 as expressed as phenotype *a_i_* and 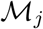 as *a_j_*, there is no inferring phenotype *a*_*i*∪*j*_ solely from phenotypes *a_i_* and *a_j_*.

We now consider the simplest case of a resident population in which a mutant appears. The mutation set of the resident population is the empty set 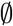, and the mutation set of the mutant is {1}, i.e., 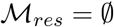 and 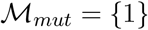. Then the following transduction events may occur:

- Interaction between a cell and a GTP carrying DNA of the same genotype: no genotypic or phenotypic change.
- Interaction between a resident cell and a GTP carrying mutant DNA: since 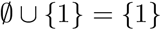, the resident cell acquires the mutation set of the mutant and changes to the mutant phenotype.
- Interaction between a mutant cell and GTP carrying resident DNA: since 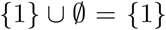, the mutation set of the mutant is unchanged and there is no phenotypic change.

Thus, whether they interact with resident or mutant bacteria, resident GTPs do not alter the phenotype of the recipient cells, while mutant GTPs may change the phenotype of resident cells into the mutant phenotype. As a consequence, the bacterial population remains dimorphic (i.e., transduction does not create a third phenotype) and the resident GTP population has no effect on the overall population dynamics.

### Phage-bacteria population dynamics with mutation and transduction

We use a multitype logistic model of birth and death (Karlin and Taylor 1981; Ethier and Kurtz 2009), a time-continuous branching process, to model the population dynamics of resident bacteria with trait value *a* and mutant bacteria with trait value *A* (population sizes *X* and *Y*, respectively), viruses (population size *Z*), and GTP carrying mutant DNA (population size *U*). The order of magnitude of the bacterial population size is denoted by *K*. Bacteria reproduce asexually at rate *b* and die at rate *d*. Both rates may depend on the population size and trait value; for simplicity, we focus on the case where the negative demographic effect of competition for limited resources is borne out by the death rate. Viruses form a homogeneous population, and GTPs are defined by the genotype and phenotype of their host of origin. Virions die at rate *d_v_*, and interactions between resident bacteria and viral particles carrying mutant DNA occur at the individual rate *η*(*a*). An interaction between a bacterium and an active virus results in instant lysis and a viral burst whereby *V* new viral particles are created, each having a probability *γ* of being a GTP. When a bacterium with trait *a* and a GTP from a cell with trait *a′* come into contact, with probability *ψ*(*a, a′*) the foreign DNA fragment is integrated by recombination into the recipient bacterial genome; the smaller the difference between *a* and *a′*, the larger *ψ*(*a, a′*) is (Touchon et al. 2017). The full mathematical derivation of the population dynamics model can be found in **Supplementary note S1**.

In the large population limit (large *K*), population sizes *X, Y, Z, U* can be rescaled into population densities *x, y, z, u* whose time dynamics are governed by the following deterministic equations (see **Supplementary note S2**):

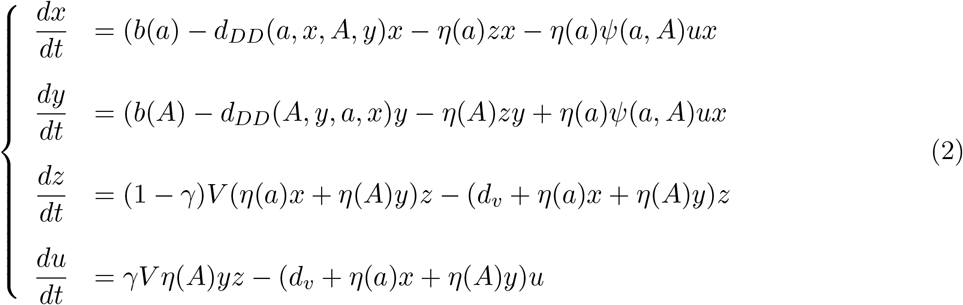

Competition among cells is assumed to result in a density-dependent (DD) death rate which increases linearly with density, given by

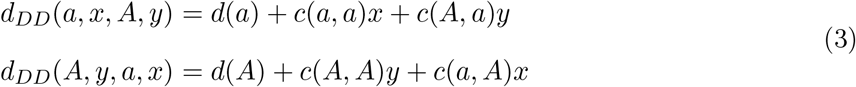

where *d*(*a*) (respectively *d*(*A*)) is the intrinsic death rate of individuals with trait value *a* (resp. *A*) and *c* measures the intensity of competition between individual cells as a function of their trait values. In a monomorphic population with trait value *a*, transduction alone has no effect on bacteria’s phenotype, hence the phage-bacteria population equilibrium:

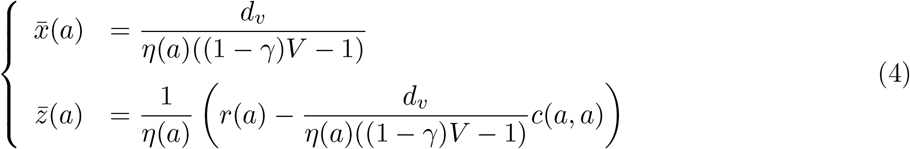

In our numerical simulations, for the chosen range of parameter values (see **Supplementary Table 1**), this equilibrium was always found to be globally stable. Virus-bacteria population cycles or the extinction of one or both populations were never detected over our range of trait values.

### Mutant invasion, Trait Substitution Sequence, and canonical equation of adaptive dynamics

We assume that the adaptation process is mutation-limited (mutation occurs rarely on the timescale of population dynamics) and that mutations have small phenotypic effects. Those are key assumptions under which the adaptation process in very large populations can be described as a *Trait Substitution Sequence* (TSS) (Metz, Geritz, et al. 1995, Champagnat 2006). In the TSS model, a mutant either invades and replaces the resident type, or goes to extinction – provided the population has not come too close to a potential evolutionary optimum (Geritz et al. 2002, Geritz 2005). This means that no two populations of resident and mutant bacteria that are phenotypically similar may coexist, except possibly near an evolutionary optimum. Thus the population evolves adaptively by making small phenotypic ‘jumps’ in the trait space, each jump corresponding to the invasion of a successful mutant into the former resident-trait population.

The invasion success of a mutant *A* in a resident population of trait value *a* is determined by the mutant’s invasion fitness, i.e. the mutant population growth rate from initially very small density in a resident population at ecological equilibrium. This is obtained from the previous dynamical system for population densities (equation 2). Here we use *S* and *p* to denote respectively the invasion fitness and invasion probability in the absence of transduction, and *S_T_* and *p_T_* to denote invasion fitness and invasion probability with transduction.

To infer the probability of invasion and characteristic invasion time while taking the population finiteness into account, we link the deterministic dynamical system describing the dynamics of large populations to the stochastic individual-level process of birth and death events driving the dynamics of small populations (initial mutant population, resident population upon complete invasion). The full mathematical treatment is presented in **Supplementary note S4**, and we summarize here the results in the absence of transduction. Once the probability *p*(*a, A*) of invasion of a mutant trait A in a resident population of trait *a* is known, we derive the ‘jump rate’ of the adaptation process with transduction. To this end, we rescale time by *K_μ_*, where *K* is the order of population size and *μ* is the mutation rate. On this new timescale, the population evolves adaptively in trait space according to the TSS model, jumping from trait *a* to trait *A* at rate

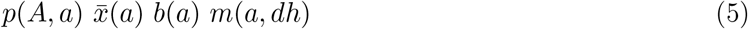

where 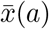 is the equilibrium population density of the resident trait *a*, and *m*(*a, dh*) is the probability of the jump from trait *a* to *a* + *h* = *A*. In this paper, the probability will only depend on the distance to resident such as *m*(*a, dh*) = *m*(*h*)*dh*: for instance, if the mutant trait is taken according to a normal law of standard deviation *σ* centered on the resident trait, we have

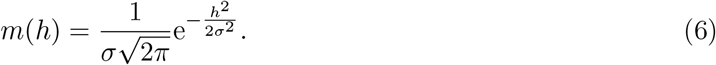

Taking the limit of arbitrarily small mutations happening fast (relative to the new timescale), the TSS converges towards a process driven by the so-called ‘canonical equation of adaptive dynamics’ (Dieckmann and Law 1996, Champagnat, Ferrière, et al. 2002, Champagnat and Méléard 2011). In the absence of transduction, this limit process is deterministic and the canonical equation reads

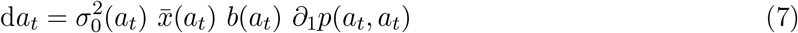

where *a_t_* denotes the trait value evolving as a function of time *t*, mutational phenotypic effects are symmetrical, and *σ*_0_ is the standard deviation of the distribution of mutational effects (mutation kernel, *m*). The direction of selection is given by the invasion probability gradient *∂*_1_*p*(*a, a*), i.e. the derivative of invasion probability *p* with respect to its first variable, evaluated for a mutant trait value equal to the resident’s, *a*. In the general case without transduction, the probability gradient can be expressed as the selection gradient *∂*_1_ *S*(*a, a*) divided by the birth rate *b*(*a*). Thus, at any time *t* the rate of adaptation is a function of the selection gradient, population size, and mutation size at that time.

‘Evolutionary singularities’ are trait values at which the selection gradient vanishes (Metz, Mylius, et al. 1996). An evolutionary singularity, denoted by *a*^*^, can be ‘attractive’ in the phenotype space, in which case the TSS of a population whose phenotypic state is some distance away from *a*^*^ will evolve under directional selection towards *a*^*^. If selection around the singularity *a*^*^ is stabilizing, the TSS remains in a neighborhood of *a*^*^. Selection may also turn disruptive around *a*^*^. Whether selection is stabilizing or disruptive is predicted by the criterion for evolutionary branching (Dieckmann and Law 1996, Champagnat and Méléard 2011).

### Simulations

We performed individual-level simulations of populations and trait dynamics by implementing a rigorous Gillespie algorithm, based on Champagnat (2006). The speed of adaptation was estimated from simulations of the TSS and compared to our mathematical approximations. All simulations were carried out using constant rates of birth, intrinsic death, and infection, respectively equal to *b*_0_, *d*_0_, *η*_0_. We used the following form for the competition term

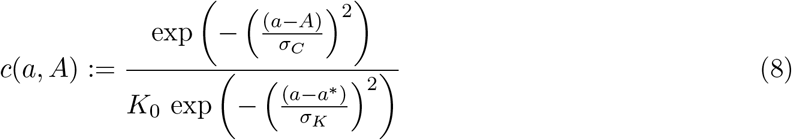

where *σ_C_* measures competition sensitivity to phenotypic difference (*a–A*), *K*_0_ represents the maximal carrying capacity of the environment, *σ_K_* represents the sensitivity of the population carrying capacity to the distance of trait *a* to the phenotypic optimum *a*^*^. We use the following form for the transduction probability

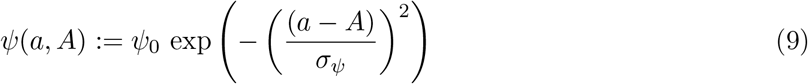

where *ψ*_0_ is the maximal transduction probability and *σ_ψ_* measures the sensitivity of *ψ* to the phenotypic difference (*a* – *A*). Default values for all parameters are listed in **Supplementary Table 1**.

## Results

### Dynamics of bacterial adaptation without transduction

First we consider the baseline scenario in which the evolving bacterial population is exposed to viral infection but no transduction occurs, *ψ*_0_ = 0. Here the invasion fitness *S*(*A, a*) of a mutant trait *A* in a resident population of trait *a* is

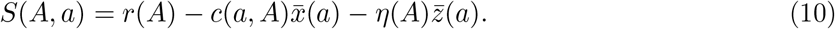

The term 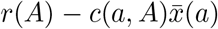 is the invasion fitness in the absence of viral infection. The additional term 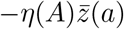 measures the negative effect of infection on bacterial fitness. As expected, this negative effect is stronger if the equilibrium viral population, 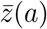, or the infection rate, *η*(*A*), is larger. From equation (10) it follows that the invasion probability is *p*(*A, a*) = [*S*(*A, a*)]_+_/*b*(*A*), with a characteristic time of invasion of the order of log(*K*) (Billiard et al. 2016).

The direction of selection is given by the selection gradient *∂*_1_*S*(*a, a*) and the rate of adaptation, by the canonical equation

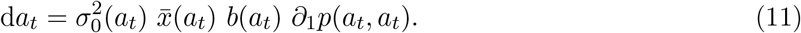

We assume that the functions *r*, *c* and *η* are such that there is a single evolutionary singularity, *a*^*^. For the model specifications used in our numerical simulations, *a*^*^ exists and is always globally attractive; selection around *a*^*^ is stabilizing if *σ_K_* < *σ_C_*, or disruptive, with evolutionary branching, if *σ_K_* > *σ_C_*. These conditions are identical to those obtained in the absence of infection (Doebeli and Dieckmann 2000).

### Mutant invasion with transduction

With transduction (*ψ*_0_ > 0), invasion fitness *S_T_*(*A, a*) of a mutant trait value *A* in a resident population of trait *a* is given by

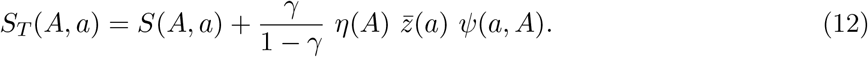

Transduction alters invasion fitness by adding the positive term 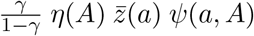. Thus, invasion fitness with transduction is always greater than invasion fitness without transduction: *S_T_*(*A, a*) > *S*(*A, a*) (Figures 2a-b). We can further decompose the additive term into three factors: 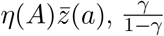, and *ψ*(*A, a*). The first factor represents the rate at which mutant bacteria interact with the viral population at ecological equilibrium with the resident bacterial population, each of these interactions being a potential opportunity for the release of GTPs. The second factor measures the abundance of GTPs released by a mutant infection relative to active viruses. Taken together, the two factors measure the strength of potential transduction events compared to infection events. The last term represents the proportion of these potential transduction events that result in actual genetic transfer. The three factors together thus quantify the intensity of transduction.

**Figure 2.**
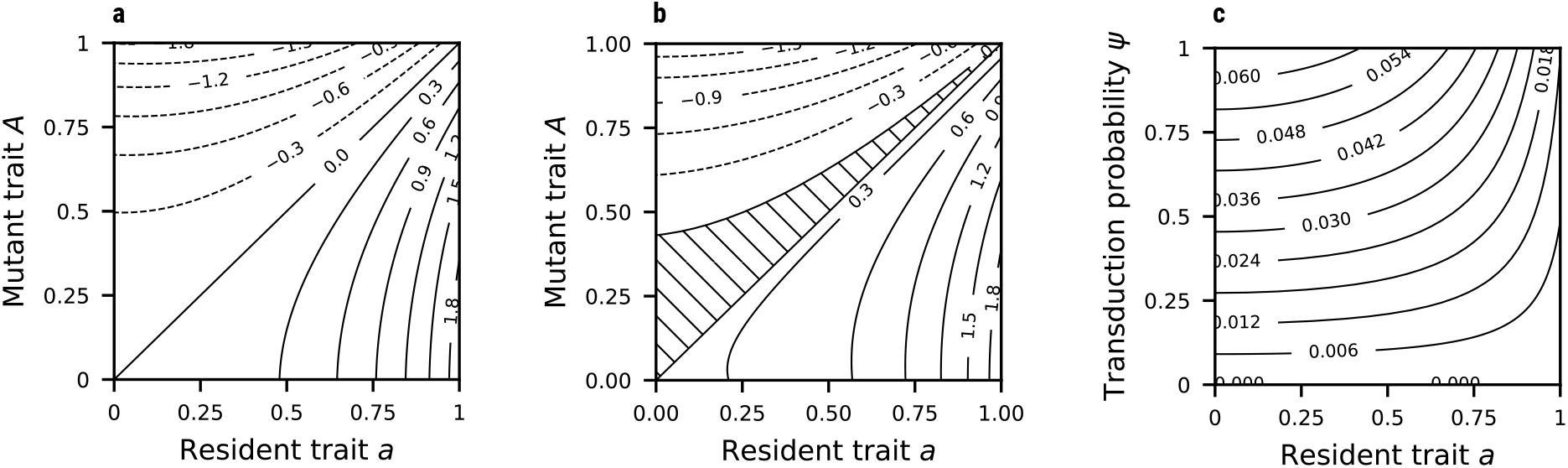
Influence of transduction on invasion fitness and the probability of invasion. **(a)** Contours of invasion fitness *S*(*A, a*) in the absence of transduction. **(b)** Contours of invasion fitness *S_T_*(*A, a*) with tranduction (*ψ*_0_ = 1.0 and *η*_0_ = 10^-5^). The hatched area highlights trait values of *a* and *A* for which *S*(*A, a*) < 0 and *S_T_* (*A, a*) > 0. **(c)** Invasion probability *p_T_* (*a, a*) of a mutant that is phenotypically identical to the resident strain, as a function of the resident trait value and transduction probability. Other parameters are set to their default values (see Supplementary Table 1).

As a consequence, mutants that are selected against (negative invasion fitness, *S* < 0) in the absence of transduction may be favored and invade when transduction is strong enough, resulting in *S_T_* > 0 (Figure 2b). Also, irrespective of the intensity of transduction, negatively selected mutants (*S* < 0) that are phenotypically close to the resident phenotype always invade. In fact, with transduction, any mutant of sufficiently small effect is predicted to invade.

Transduction also affects the mutant-resident dynamics following mutant invasion. The potential for ‘back invasion’ by the resident population once rare is determined by the ‘back invasion fitness’ (see **Supplementary note S2**)

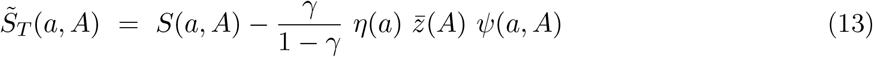

By making a negative contribution to the right-hand side of equation (13), transduction generally hampers back invasion. In particular, for any mutation of small effect (*S*(*A, a*) ≈ 0), transduction always begets invasion 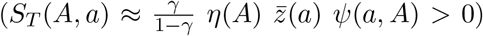 and always prevents back invasion 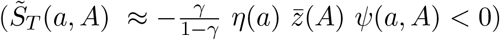.

In **Supplementary note S3** we show that transduction always increases the probability of invasion of a mutant. We also show that for small transduction rates (which is the case when mutations have small phenotypic effects) and large viral burst sizes (*V* >> 1, which is expected in natural systems), the probability of invasion with transduction is

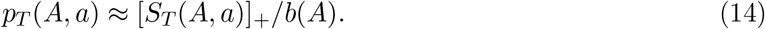

and the characteristic time of invasion remains of order log(*K*).

Of note, equations (12) and (14) imply *S_T_*(*a, a*) > 0 and *p_T_*(*a, a*) > 0 for any trait value *a*. Thus, a mutant that is phenotypically identical to the resident type has positive invasion fitness (Figure 2b), as opposed to zero invasion fitness in models without transduction (Figure 2a). This is revealed by our explicit genotype-phenotype map: albeit phenotypically identical to the resident type, the mutant type is genetically different (by definition, it carries a new mutation) and our model of transduction accounts for this. Using notations introduced earlier, we have 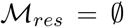 and 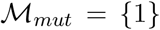, and our model predicts 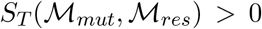. With this modeling framework, phenotypically similar strains can have different invasion fitnesses, and only when two strains are genotypically equivalent 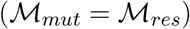 that the invasion fitness is equal to zero.

The invasion probability for mutants that are phenotypically identical to their resident progenitor, *p_T_*(*a, a*), provides an approximation for the invasion probability of mutations of small phenotypic effect. The probability *p_T_*(*a, a*) is higher near the evolutionary singularity *a*^*^ (Figure 2c). Thus, due to transduction, the invasion probability of any mutation of small effect increases as the adaptation process brings the population closer to the evolutionary singularity.

### Rate of adaptation with transduction

Here we assume that mutations have small phenotypic effects. By rescaling time appropriately, we derive a macroscopic model for the dynamics of adaptation similar to the canonical equation. With transduction, however, the macroscopic model retains a stochastic component and takes the form of a stochastic differential equation or integro-differential equation, depending on the shape of the mutation kernel. In **Supplementary note S4**, we show that the models, hence the dynamics of adaptation that they capture, critically depend on whether the mutation kernel is symmetrical around zero (unbiased mutational effects) or not (biased mutational effects).

#### Case of unbiased mutation

With unbiased mutation the rate of adaptation is controlled by

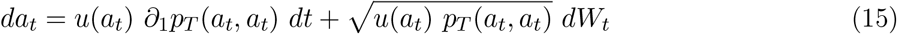

where 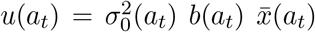 and *W_t_* is a Brownian motion. The Brownian component arises because transduction causes *p_T_*(*a, a*) > 0. The mean adaptation rate with transduction is given by the deterministic part of the stochastic canonical equation (15). To assess the effect of transduction on adaptation speed away from the evolutionary optimum, we compare this deterministic part to the similar term in the canonical equation without transduction, equation (11). Under our simulation assumptions (i.e. birth rate, infection rate, and mutation size independent of the trait), *u*(*a_t_*) is constant, and the rate of adaptation is solely determined by the invasion probability *p_T_*(*a_t_, a_t_*) and its first derivative *∂*_0_*p_T_*(*a_t_, a_t_*).

The adaptation rate is always higher with transduction, all the more so as the maximum transduction rate, *ψ*_0_, is larger (Figures 3a-c). As evolutionary trajectories come closer to the optimum, the (absolute) difference in adaptation rate with vs. without transduction becomes smaller (Figure 3b), but this effect is largely driven by the general tendency for adaptation to slow down near the optimum. When controlling for the slow down by computing the relative difference in adaptation rates, the accelerating effect of transduction remains (Figure 3c). Numerical simulations of the TSS agree with these analytical results (Figures 3d-f).

**Figure 3.**
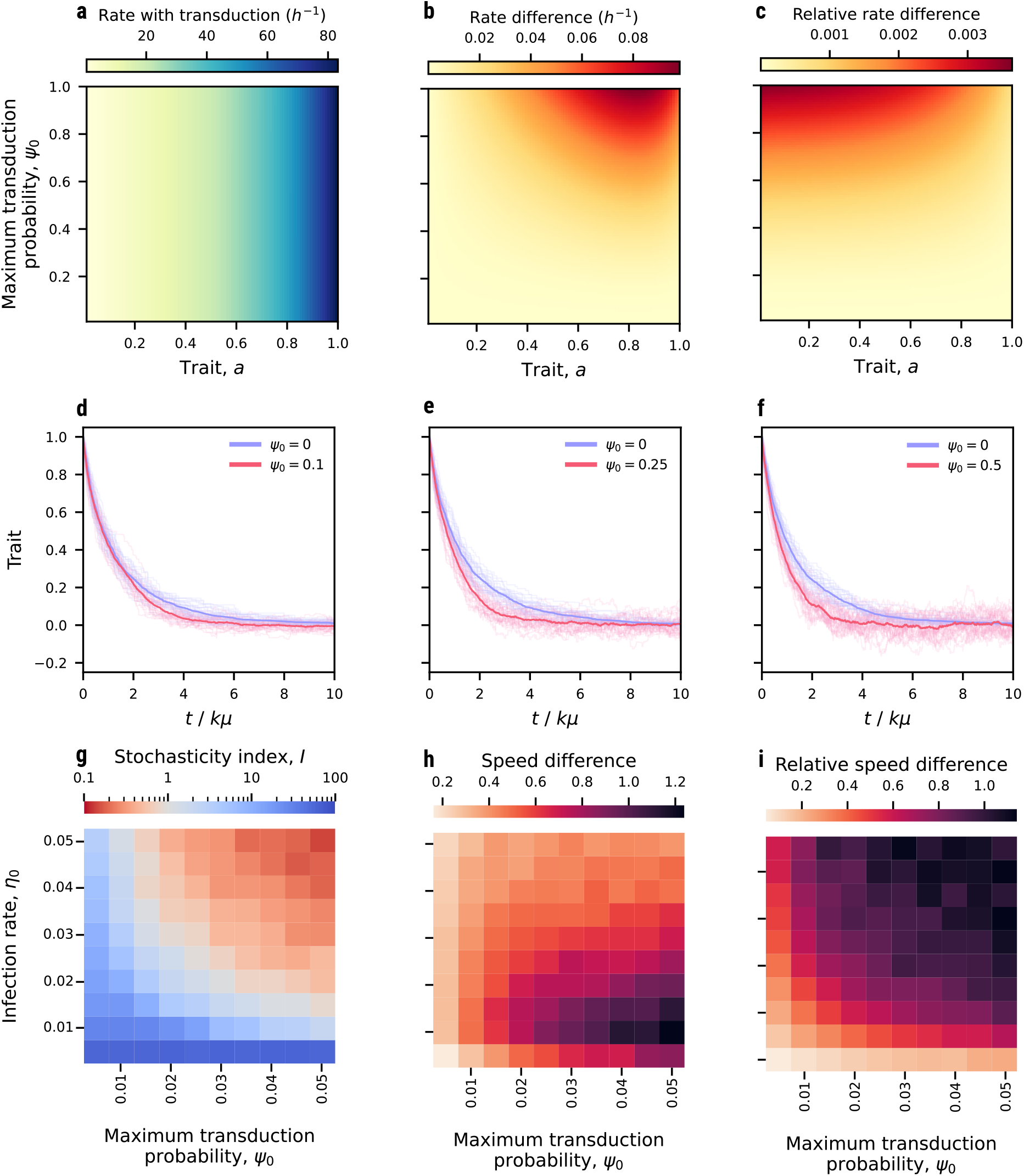
Influence of transduction on adaptation rates. (a) Rate of adaptation with transduction as a function of trait value *a*, for a given maximum transduction probability, *ψ*_0_. (b) Absolute and (c) relative difference between rates of adaptation with and without transduction. The evolutionary optimum is at *a*^*^ = 0. (d-f) Simulated Trait Substitution Sequences (TSS) for *η*_0_ = 10^-5^ and (d) *ψ*_0_ = 0.01, (e) *ψ*_0_ = 0.25, and (f) *ψ*_0_ = 0.5. 30 simulation runs for each *ψ*_0_ value were performed (light blue and light red lines) with a time normalization by a factor of *K_μ_*. The thick dark lines represent the mean trajectory for each set of simulations. (g) Adaptation index, *I*. Blue and red shades respectively indicate *I* > 1 (the dominant effect of transduction is to accelerate adaptation away from the optimum) and *I* < 1 (the dominant effect of transduction is stochasticity around the optimum). (h) Absolute and (i) relative difference in convergence adaptation speed, *α*, with vs. without transduction, calculated from 100 simulation runs for each pair of transduction probability and infection rate. Other parameters are set to their default values (see Supplementary Table 1). See text for definitions of *I* and *α*.

Because the invasion probability *p_T_*(*a, a*) increases as trait *a* approaches *a*^*^ (Figures 2c), we expect the dynamics of adaptation to become more stochastic as the population evolves closer to the singularity *a*^*^. This is confirmed by numerical simulations of the TSS (Figures 3d-f). Thus, transduction increases both adaptation speed away from the optimum, and stochasticity near the optimum. To quantify the net effect of these antagonistic influences on adaptation, we use an index of adaptation, *I*, defined as the ratio of the speed of adaptation without transduction relative to the mean sojourn time of the population in a small phenotypic neighborhood around the evolutionary optimum (see **Supplementary note S5** for more detail). *I* ≪ 1 means that the stochastic component of adaptation pushes the process away from the evolutionary optimum faster than the deterministic driver can pull it back. In this case, the population tends to remain in a maladapted state, i.e. away from the predicted optimum. Conversely, *I* ≫ 1 means that the deterministic driver of adaptation brings the population near its evolutionary optimum faster than the stochastic component upsets adaptation. In this case, adaptation trajectories can be approximated with exponential functions of time *t* of the form *C* exp(–*αt*). The parameter *α* then provides a measure of convergence speed that can be compared across models with and without transduction.

We used numerical simulations to evaluate how the adaptation index, *I*, and adaptation convergence speed, *α*, respond to two key parameters of the bacteria-phage interaction, the rate of infection, *η*_0_, and the probability of transduction, *ψ*_0_ (Figures 3g-i). If either one of the infection rate, *η*_0_, or transduction probability, *ψ*_0_, is very low (less than ca. 0.01), then *I* ≫ 1 and the adaptation trait dynamics closely follows its predicted deterministic path towards *a*^*^ (Figures 3d,g). If both *η*_0_ and *ψ*_0_ are large enough (product larger than ca. 5 10^-4^), then *I* ≪ 1, meaning that even though convergence towards an evolutionary optimum is expected, the adaptation trajectories strongly fluctuate around the optimum without ever settling down (Figures 3e-g). Increasing the transduction probability generally increases the adaptation convergence speed (Figures 3d-f,h). The absolute effect, in comparison with the convergence speed without transduction, is sensitive to the infection rate, being strongest at low infection rates (Figure 3h). This is because the mortality cost of infection tends to attenuate the accelerating effect of transduction on adaptation. When measuring the effect of transduction on the convergence speed relatively to the speed without transduction (*relative* convergence speed, Figure 3i), we find that the larger the infection rate, the stronger the positive effect of transduction on the speed of adaptation.

#### Case of biased mutation

With biased mutation, the average mutational effect is nonzero and given by

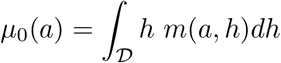

Then the characteristic timescale over which adaptation proceeds is set by *t/ϵ*, where *t* is the baseline timescale over which the cells’ population dynamics unfold. The *t/ϵ* timescale is much shorter than the *t/ϵ*^2^ timescale over which adaptation proceeds with unbiased mutation, i.e. adaptation occurs much more rapidly with biased mutation. On this faster timescale, the stochasticity of the adaptation process with transduction is smoothed out, and the adaptation dynamics are governed by a deterministic ordinary equation

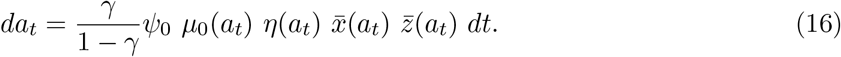

In this case, the long-term adaptation dynamics takes steps of the order of the average mutational bias, measured by *μ*_0_(*a*), at a rate given by the population rate of transduction events, equal to 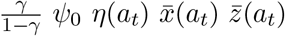.

### Effect of transduction on bacterial adaptive diversification

Without transduction, the model predicts conditions under which selection turns from directional to disruptive around the evolutionary singularity *a*^*^, resulting in evolutionary branching (Figure 4a). A full mathematical investigation of evolutionary branching with transduction goes beyond the scope of this paper. Instead we gained some insights into the effect of transduction on bacterial adaptive diversification from a large set of numerical simulations. The simulations consistently show that even the lowest level of transduction widens the trait range over which a mutant can invade in a neighborhood of the evolutionary singularity (Figures 4b-f). None-the-less, evolutionary branching occurs under the conditions predicted in the absence of transduction, and the fact that back invasion fitness is always negative does not seem to slow the process.

**Figure 4.**
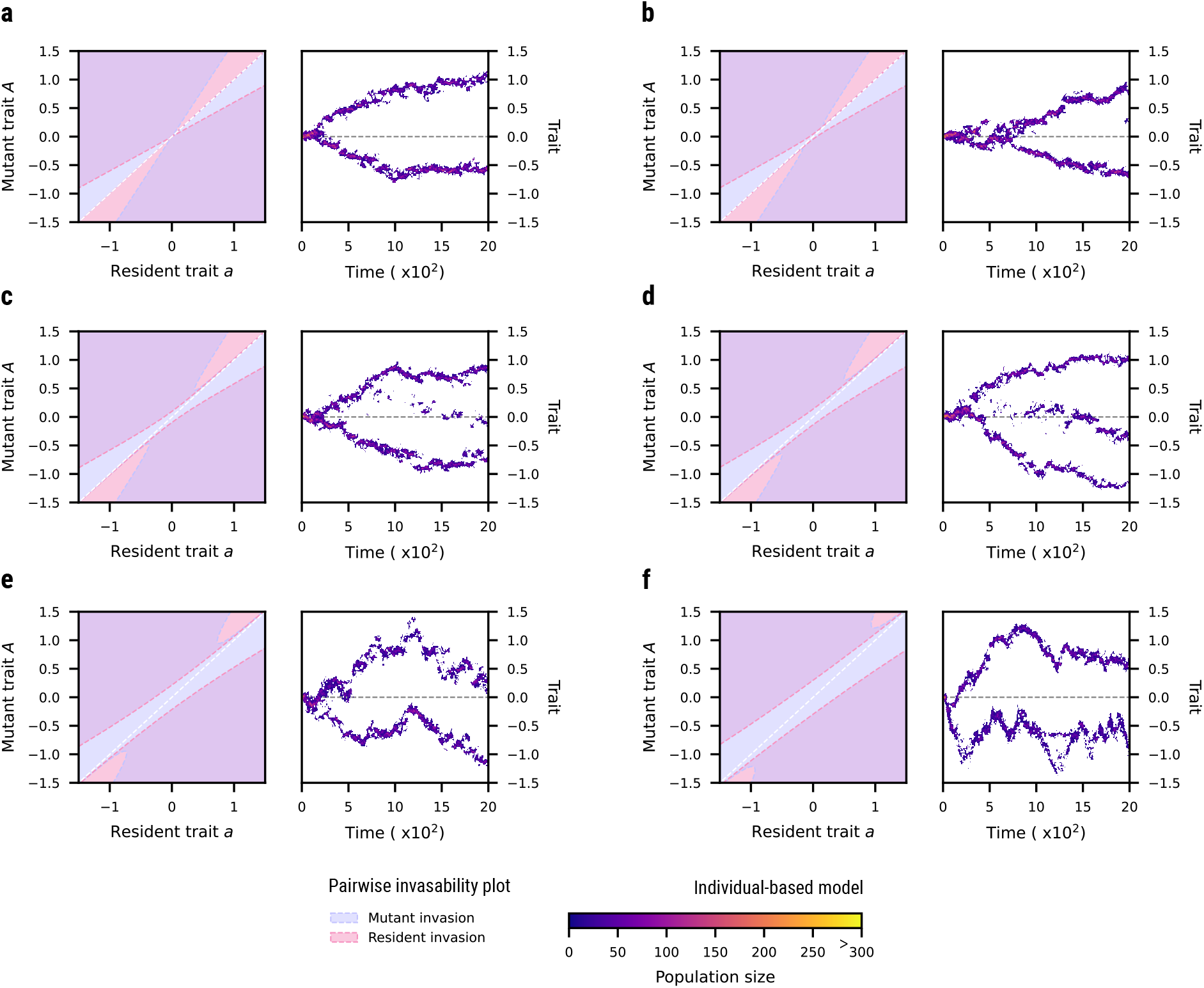
Pairwise invasibility plots and example simulations of trait distribution dynamics. The transduction probability, *ψ*_0_, takes the following values: (a) 0; (b) 0.01; (c) 0.05; (d) 0.1; (e) 0.25; (f) 0.5. For each value of *ψ*_0_, the left pane shows regions of resident and mutant trait values for which mutant invasion fitness is positive (blue) or resident invasion fitness is positive (red). In the region of overlap (purple), the mutant and resident populations may coexist. Right panes show trait distribution dynamics simulated by making use of a Gillespie algorithm. Parameter values: *η*_0_ = 10^-5^, *σ_K_* = 2, *σ_C_* = 1, and other parameters set to their default values (see Supplementary Table 1).

At low transduction rate (low *ψ*_0_), a third, intermediate branch can evolve and station around the evolutionary singularity, which is a fitness minimum (Figures 4b-d). This branch emerges from the mixing of genomes in the two outer branches (as explained in Figure 1) and follows either one of these branches. The new branch persists only due to horizontal gene transfer, and would go extinct from competition with the external branch without the flux induced by transduction. Thus, competition exerted by individuals in the intermediate branch select for further divergence of the outer branches (Figure 4d). In sum, transduction at low rates tends to increase the genetic and phenotypic diversity and phenotypic range that evolves in the population.

At higher transduction rates, large stochastic variations manifest in the two outer branches. Large stochastic excursions of one branch towards the evolutionary singularity tend to outcompete any middle branch (Figures 4e,f). Large stochastic excursions of one branch away from the singularity may allow the other branch to wander around the singularity (Figures 4e,f). Thus, transduction makes it possible for a subpopulation to station around an evolutionary optimum which is a fitness minimum. When this happens, competition exerted by this sub-population prevents any third branch from even sprouting (Figures 4e,f). See Supplementary figure 1 for additional example simulations.

## Discussion

Horizontal gene transfer is expected to have conflicting effects on bacterial adaptation: the transfer of beneficial mutations may facilitate and accelerate adaptation, while the transfer of deleterious mutations might hold adaptation back. Transduction by virulent phages adds a direct demographic factor to these conflicting effects, since the frequency of gene transfer is tied to viral infection hence to cell mortality. To resolve the effect of transduction on bacterial adaptation, we extended the trait-based approach of adaptive dynamics modeling to include a simple genotype-phenotype map and capture the mobility of genetic elements between host cells and their potential phenotypic effects. Our hybrid – genetic and trait-based – approach uses an infinite-site, infinite-allele model with small additive effects of mutations. Our model applies to generalized transduction by virulent phages and assumes that ‘back transduction’ of a descendant allele recombining at the progenitor locus is a very unlikely event, which is consistent with the non-homologous end joining transduction mechanism (Popa, Hazkani-Covo, et al. 2011).

In the absence of transduction, the mortality cost of viral infection tends to slow bacterial adaptation down. A key effect of transduction is to increase the invasion fitness of any mutant, irrespective of their fitness effect (deleterious or beneficial) in the absence of transduction. This makes it possible for deleterious alleles to invade, and for beneficial alleles to invade faster. Far from an evolutionary optimum, the positive fitness effect on deleterious mutations (of a given mutational effect size) is small relative to the effect on beneficial mutations. As a consequence, adaptation can be much faster with transduction than without. Close to an evolutionary optimum, transduction tends to reduce the fitness difference between deleterious and beneficial mutants and both can have relatively high positive invasion fitness. This can generate strong stochastic fluctuations in the adaptive trajectories. If the evolutionary optimum is a fitness minimum in the absence of transduction, thus turning selection from directional to disruptive, the expected phenotypic polymorphism and divergence also occurs with transduction. At low transduction rates, a subpopulation may even persist near the optimum, driving further phenotypic divergence of the other coexisting subpopulations. At high transduction rates, stochasticity drives large fluctuations in phenotypic branches, including recurrent incursions near the optimum.

Previous studies (e.g. Moradigaravand and Engelstädter 2014) have modeled and discussed the ‘bad gene effect’ of HGT. Non-homologous end joining transduction is distinct in the sense that the bad gene effect never consistently reverts the direction of selection, as it may in the case of conjugation (Billiard et al. 2016). Rather, it tends to broaden the range of selection outcomes. As the transfer term of invasion fitness *S_T_*(*A, a*) is positive for any mutant trait that arises in the resident population, not only may deleterious trait values invade, but beneficial trait values have even greater invasion fitness than they would in the absence of transduction, implying a larger probability of invasion and a shorter invasion time. Quantitatively, these effects can be substantial (Figure 2b). This prediction is consistent with previous theoretical results on HGT (Cohen et al. 2006) and experimental data showing that horizontal gene transfer can facilitate adaptation by accelerating the fixation of beneficial alleles (Baltrus et al. 2008).

With transduction, any small phenotypical variant has positive invasion fitness. As a consequence, even though directionality towards the evolutionary optimum remains favored (as shown by the deterministic term in the canonical equation (15)), random jumps of the TSS in the opposite direction are possible at any mutation event. This broad range of invading mutants around the evolutionary optimum fuels wide stochastic fluctuations in the adaptation process. Stochastic fluctuations of the adaptation trajectory due to transduction are most pronounced near the evolutionary optimum, *a*^*^, thereby hampering convergence to the evolutionary optimum and preventing the stabilization of adaptive trajectories around it. This is a consequence of the ‘top-down’ control of the bacterial population by viruses, which means that for traits closer to the evolutionary optimum, the population of viruses (not bacteria) becomes larger, resulting in a higher rate of transduction and eventually a wider range of mutant phenotypes with positive invasion fitness around *a*^*^.

Stochasticity in adaptive evolution is generally associated with genetic drift in finite (small) population size. In contrast, the adaptation stochasticity evidenced here occurs in spite of the large population size, and its long-term effect on the adaptive trait dynamics is captured by the stochastic canonical equation (15). This equation can be compared with the canonical equation for finite populations (Champagnat, Lambert, et al. 2007). Both equations contain a stochastic term proportional to the square root of the invasion probability of a neutral trait. Classically, the non-zero value of this term reflects the effect of genetic drift in finite populations. Here, the population is virtually infinite and the non-zero value comes from the stochasticity of transduction events. Although the root cause of stochasticity is profoundly different, the outcome is qualitatively similar: deleterious alleles can invade and go to fixation.

The fact that trait fluctuations still occur near the evolutionary optimum implies that the host population will never be fully adapted to a static environment: by facilitating the fixation of *any* mutant, transduction lowers the mean fitness of the population. This is in line with previous studies showing that HGT is not a facilitator of adaptation in static environments (Raz and Tannenbaum 2010). In contrast, transduction is expected to accelerate adaptation in response to environmental change that changes the evolutionary optimum, all the more so as the shift in optimum is wider.

Our results also shed new light on the effect of horizontal gene transfer on genetic and phenotypic diversity (Niehus et al. 2015). Transduction events add new lineages to the population (Figure 1), with two contrasting outcomes that are revealed by individual-level simulations (Supplementary figure 1), either enhanced diversification or transient optimization. At low transduction rates, the genome-mixing effect of transduction dominates and trait diversity increases. At higher transduction rates, the stochasticity of transduction events scale up and cause the stochastic motion of evolutionary branches; this may result in ‘transient optimization’ whereby one branch (sub-population) stations close the evolutionary optimum, thus causing extinction of some or all the other branches, hence a transient reduction in trait diversity.

Our analysis highlights the influence of transduction rate parameters on the rate and dynamics of bacterial adaptation. These parameters are: the infection rate, *η*, the gene transducing particles (GTP) production fraction, *γ*, and the maximum transduction probability, ψ_0_. The latter two, that are specific to the transduction process, remain poorly constrained. Laurenceau et al. (2021) provided evidence suggesting that generalized transduction is the dominant mode of transduction in *Prochlorococcus*, the most abundant phototrophs in the open ocean. By examining the mispackaging of host DNA by different cyanophages, they described a mechanism to explain the production of GTPs, which paves the way toward quantifying GTP production. Interestingly, their proposed mechanism suggests that GTP production might respond to environmental conditions and vary with ocean depth, latitude, and season. Quantitative data are also lacking for the maximum transduction probability, ψ_0_. It is known that multiple barriers exist to prevent the recombination of exogenous DNA in the recipient cell (Popa and Dagan 2011; Oliveira et al. 2016), and here too, a better understanding of the mechanism should help estimating this critical parameter. The development of new sequencing-based methodology, e.g. Kleiner et al. (2020) transductomics approach, will inform these efforts by providing insights into real-time ongoing HGT in complex microbial communities.

Our model is focused on generalized transduction by virulent phages. Future developments will aim at extending the model to temperate (lysogenic) phages, which can also drive specialized and lateral transduction. This will broaden the scope of our framework to address questions involving the evolution of transduction-related traits in both bacterial host and phage populations. All three transduction parameters may be under the genetic control of both phage and host – the infection rate by virulent genes and resistant genes, respectively; the GTP production rate and transduction probability by viral genes that control DNA excision and packaging and host genes that control DNA integration. With temperate phages that can transfer bacterial DNA without killing recipient cells, positive selection pressures may exist on genes that benefit both hosts and phages (Fillol-Salom et al. 2019; Wendling et al. 2021) and result in high-frequency transduction (Chen et al. 2018).

For transduction involves the transfer and integration of alleles between hosts, models that keep track of the individuals’ phenotype (but not genotype) are insufficient to describe the adaptive trait dynamics of a host population in which transduction occurs. Our modeling approach goes beyond previous population-genetic models of HGT (Niehus et al. 2015, Mao and Lu 2016, Nazarian et al. 2018) to track the genealogy of mutations together with phenotypic trait values, which influence demographic performance and ecological interactions among individuals and their environment. Beyond the question of how transduction affect host adaptation, the mathematical method used here could pave the way to further advance the integration of transmission genetics in trait-based models of adaptation.

## Author contribution

R.F. and S.M. conceived the study. All authors developed the model. P.C. performed the analysis. All authors wrote the first version of the manuscript.

## Acknowledgements

This work is supported by France Investissements d’Avenir program (ANR-10-LABX-54 MemoLife, ANR-10-IDEX-0001-02 PSL), PSL - University of Arizona Mobility Program and the Chair “Modélisation Mathématique et Biodiversité” of Veolia Environnement-Ecole Polytechnique-Museum National d’Histoire Naturelle-Fondation X. P.C. is supported by a doctoral fellowship from the French IPEF program. R.F. acknowledges support from the U.S. National Science Foundation, Dimensions of Biodiversity (DEB-1831493), Biology Integration Institute-Implementation (DBI-2022070), and National Research Traineeship (DGE-2022055) programs; and from the United States National Aeronautics and Space Administration, Interdisciplinary Consortium for Astrobiology Research program.

## Data accessibility

no data is used in this paper.

