## Supplementary information for "Viral transduction and the dynamics of bacterial adaptation"

### Supplementary note S1: A stochastic model for mutant invasion with transduction

We consider a population of five types of individuals, under the hypothesis that initial resident population sizes are of order $K$, which will be used to scale parameters when we study the case of large population ($K \to \infty$). The trait value of resident bacteria is $a$ (with the $\emptyset$ mutation set) while the trait value of mutant bacteria and gene transducing particles (GTP) is $A$ (with the $\{1\}$ mutation set). We call $\mathcal{D} \subset \mathbb{R}$ the domain of feasible trait values; we thus have $a, A \in \mathcal{D}$. The four populations involved in the mutant invasion process are

- The resident population, adding to $N_t^{a,K}$ individuals with trait $a$ at time $t$
- The mutant population, adding to $N_t^{A,K}$ individuals with trait $A$ at time $t$
- The viruses found in the environment, adding to $V_t^K$ individuals at time $t$
- The GTPs coming from the mutant population found in the environment, adding to $G_t^K$ individuals at time $t$

Note that GTPs coming from the resident population do exist, but they have no ecological impact. We define population densities at time $t \geq 0$ for the different types of individuals as

$$(X_t^K, Y_t^K, Z_t^K, U_t^K) = \frac{1}{K}(N_t^{a,K}, N_t^{A,K}, V_t^K, G_t^K). \tag{1}$$

The population changes in continuous time, undergoing events of birth, death, lysis due to infection, or transduction. For simplicity, we assume that lysis is instantaneous once a bacterial cell is infected. For $u, v \in \{a, A\}$ we define the following:

- $b_K(u)$ is the rate at which an individual with trait $u$ gives birth to a new individual with trait $u$.
- $d_K(u)$ is the intrinsic rate at which an individual with trait $u$ dies.
- $c_K(u, v)$ for $v \in \{a, A\}$ measures the effect of competition exerted by an individual with trait $v$ on an individual with trait $u$.

- $\eta_K(u)$ is the rate at which an individual with trait $u$ encounters a viral particle, be it a virus or a GTP.

- $V$ is the number of viral particles released during a lysis event. Each of these $V$ viral particles, independently of one another, has an intrinsic probability $\gamma$ to be a GTP and a probability $1-\gamma$ to be a virus. Therefore, we define $p_i^{(V,\gamma)}$ for $i \in \{1, ..., V\}$ the probability that $i$ particles of the burst are GTPs. $(p_i^{(V,\gamma)})_i$ follows a binomial distribution of parameter $(V, \gamma)$.

- $\psi(u, v)$ for $u, v \in \{a, A\}$ is the probability that the trait $v$ is successfully transferred from a GTP to a bacterial cell with trait $u$ during a transduction event. Typically, the closer two traits $a, A$ are, the great the chance the DNA strand be transferred, so $\psi$ is taken of the form $\psi(a, A) := \psi(||a - A||)$ where $||.||$ denotes a distance measure and $\psi$ decreases with distance.

- $d_v$ is the intrinsic rate at which a viral particle dies.

We assume that all these parameters are uniformly bounded with respect to $K$, respectively by $\bar{b}, \bar{d}, \bar{c}, \bar{\eta}$.

We know that the process $(X_t^K, Y_t^K, Z_t^K, U_t^K)_{t\geq 0}$ is fully described by its infinitesimal generator applied to continuous bounded test functions $f$ from $\mathbb{R}^4$ to $\mathbb{R}$. For $(x, y, z, u) \in (\mathbb{N}/K)^4$:

$$
\begin{aligned}
L^K f(x,y,z,u) =& K b_K(a) x \left( f(x + \frac{1}{K}, y, z, u) - f(x,y,z,u) \right) \\
&+ K b_K(A) y \left( f(x, y + \frac{1}{K}, z, u) - f(x,y,z,u) \right) \\
&+ K(d_K(a) + K c_K(a,a) x + K c_K(a,A) y) x \\
&\times \left( f(x - \frac{1}{K}, y, z, u) - f(x,y,z,u) \right) \\
&+ K(d_K(A) + K c_K(A,a) x + K c_K(A,A) y) y \\
&\times \left( f(x, y - \frac{1}{K}, z, u) - f(x,y,z,u) \right)
\end{aligned}
$$

$$
\begin{aligned}
&+K^2\eta_K(a)xz\sum_{i=0}^{V}p_i^{(V,\gamma)}\\
&\quad\times\left(f(x-\frac{1}{K},y,z+\frac{V-i-1}{K},u)-f(x,y,z,u)\right)\\
&+K^2\eta_K(A)yz\sum_{i=0}^{V}p_i^{(V,\gamma)}\\
&\quad\times\left(f(x,y-\frac{1}{K},z+\frac{V-i-1}{K},u+\frac{i}{K})-f(x,y,z,u)\right)\\
&+K^2\eta_K(A)yu\\
&\quad\times\left(f(x,y,z,u-\frac{1}{K})-f(x,y,z,u)\right)\\
&+K^2\eta_K(a)xu\psi(a,A)\\
&\quad\times\left(f(x-\frac{1}{K},y+\frac{1}{K},z,u-\frac{1}{K})-f(x,y,z,u)\right)\\
&+K^2\eta_K(a)xu(1-\psi(a,A))\\
&\quad\times\left(f(x,y,z,u-\frac{1}{K})-f(x,y,z,u)\right)\\
&+KVd_{v,K}z\left(f(x,y,z-\frac{1}{K},u)-f(x,y,z,u)\right)\\
&+KVd_{v,K}u\left(f(x,y,z,u-\frac{1}{K})-f(x,y,z,u)\right).
\end{aligned}
$$

Let us comment on this generator row by row (we omit certain rows due to the parallel roles individuals with trait $a$ and $A$ play in it):

$$Kb_K(a)x\left(f(x+\frac{1}{K},y,z,u)-f(x,y,z,u)\right)$$

represents the birth event of a new individual with trait $a$.

$$K(d_K(a)+Kc_K(a,a)x+Kc_K(a,A)y)x\left(f(x-\frac{1}{K},y,z,u)-f(x,y,z,u)\right)$$

represents the death event of an individual with trait $a$, which may be due to competition or intrinsic causes.

$$K^2\eta_K(a)xz\sum_{i=0}^{V}p_i^{(V,\gamma)}\left(f(x-\frac{1}{K},y,z+\frac{V-i-1}{K},u)-f(x,y,z,u)\right)$$

represents the infection and lysis of an individual with trait $a$.

$$K^2\eta_K(A)yz\sum_{i=0}^{V}p_i^{(V,\gamma)}\left(f(x,y-\frac{1}{K},z+\frac{V-i-1}{K},u+\frac{i}{K})-f(x,y,z,u)\right)$$

represents the infection and lysis of an individual with trait $A$. This lysis event may produce GTPs.

$$K^2\eta_K(A)yu\left(f(x,y,z,u-\frac{1}{K})-f(x,y,z,u)\right)$$

represents the interaction between a mutant bacteria and a GTP. Nothing occurs for the bacterial cell, while the GTP is lost from the population.

$$K^2\eta_K(a)xu\psi(a,A)\left(f(x-\frac{1}{K},y+\frac{1}{K},z,u-\frac{1}{K})-f(x,y,z,u)\right)$$

represents a successful transduction event.

$$K^2\eta_K(a)xu(1-\psi(a,A))\left(f(x,y,z,u-\frac{1}{K})-f(x,y,z,u)\right)$$

represents a failed transduction event.

$$Kd_{v,K}z\left(f(x,y,z-\frac{1}{K},u)-f(x,y,z,u)\right)$$

represents the death of a virion.

$$Kd_{v,K}u\left(f(x,y,z,u-\frac{1}{K})-f(x,y,z,u)\right)$$

represents the death of a GTP.

### Supplementary note S2: Large population limit and the effect of transduction on adaptation dynamics

#### Large population limit

Here we establish a deterministic approximation of the stochastic process, on the ecological timescale. In doing so, we show mathematically that for small mutational effects, less fit mutants (lower invasion fitness in the absence of transduction) can invade their resident population.

We used the following scalings. First, we define the initial population sizes as population densities:

$$\begin{aligned}
(X_0^K, Y_0^K, Z_0^K, U_0^K) &\underset{K\to+\infty}{\overset{\mathbb{P}}{\to}} (x_0, y_0, z_0, u_0) \in \mathbb{R}^4, \\
\mathbb{E}[\left(X_0^K\right)^3] &< +\infty, \\
\mathbb{E}[\left(Y_0^K\right)^3] &< +\infty, \\
\mathbb{E}[\left(Z_0^K\right)^3] &< +\infty, \\
\mathbb{E}[\left(U_0^K\right)^3] &< +\infty.
\end{aligned}$$

We rescale the individual parameters as follows when $K \to +\infty$: $\forall u, v \in \{a, A\}, b_K(u) \to b(u), d_K(u) \to d(u), Kc_K(u,v) \to c(u,v), KV\eta_K(u) \to \eta(u), d_{v,K} \to d_v$. Under this scaling, we show that the process converges in law to the unique solution of the following system of ordinary differential equations (ODEs):

$$\begin{cases}
\frac{dx}{dt} &= (b(a) - d(a) - c(a,a)x - c(A,a)y)x - \eta(a)zx - \eta(a)\psi(a,A)ux \\
\frac{dy}{dt} &= (b(A) - d(A) - c(A,A)y - c(a,A)x)y - \eta(A)zy + \eta(a)\psi(a,A)ux \\
\frac{dz}{dt} &= (1-\gamma)V(\eta(a)x + \eta(A)y)z - (d_v + \eta(a)x + \eta(A)y)z \\
\frac{du}{dt} &= \gamma V\eta(A)yz - (d_v + \eta(a)x + \eta(A)y)u.
\end{cases}$$

*Proof.* The proof follows the classical workflow that can be found in Méléard and Bansaye (2015), based on uniqueness and compactness arguments. The fact that the process involves four different types of individuals does not cause any particular difficulty. □

### Study of the dynamical system

We now turn to the study of the dynamical system

$$\begin{cases} \frac{dx}{dt} &= (b(a) - d(a) - c(a,a)x - c(A,a)y)x - \eta(a)zx - \eta(a)\psi(a,A)ux \\ \frac{dy}{dt} &= (b(A) - d(A) - c(A,A)y - c(a,A)x)y - \eta(A)zy + \eta(a)\psi(a,A)ux \\ \frac{dz}{dt} &= (1-\gamma)V(\eta(a)x + \eta(A)y)z - (d_v + \eta(a)x + \eta(A)y)z \\ \frac{du}{dt} &= \gamma V \eta(A)yz - (d_v + \eta(a)x + \eta(A)y)u. \end{cases} \tag{2}$$

We consider an invasion event of a mutant population of type $A$ in a resident population of type $a$. We first examine the ecological equilibrium between resident bacteria $\bar{x}(a)$ and viruses $\bar{z}(a)$[1] (i.e. a $(\bar{x}(a), 0, \bar{z}(a), 0)$ equilibrium) before imposing a small disturbance $(y, u)$ to the system. Before the mutant arises, the following dynamical system applies:

$$\begin{cases} \frac{dx}{dt} &= (b(a) - d(a) - c(a,a)x - \eta(a)z)x \\ \frac{dz}{dt} &= (((1-\gamma)V - 1)\eta(a)x - d_v)z. \end{cases} \tag{3}$$

We are interested in the state where both bacterial and viral populations coexist at an ecological equilibrium when the mutant bacterial strain arises. We work under the assumption that no limit cycles occur. Numerically, this is generally the case when the infection rate $\eta$ is one order of magnitude smaller than the competition coefficient $c$. We focus on equilibrium points $(\bar{x}(a), \bar{z}(a))$ with $\bar{x}(a) \geq 0, \bar{z}(a) \geq 0$. Solving (3) for equilibrium points in the first quadrant yields

$$\begin{cases} \bar{x}(a) &= \min\left(\frac{d_v}{\eta(a)((1-\gamma)V-1)}, \frac{r(a)}{c(a,a)}\right) \\ \bar{z}(a) &= \max\left(0, \frac{1}{\eta(a)}\left(r(a) - \frac{d_v}{\eta(a)((1-\gamma)V-1)}c(a,a)\right)\right) \end{cases} \tag{4}$$

where $r(a) = b(a) - d(a)$. The existence of a solution where $\bar{z}(a) > 0$ requires

[1] We write the dependencies to the trait $a$ as they will be useful later

$$0 < \frac{d_v}{\eta(a)((1-\gamma)V-1)} < \frac{r(a)}{c(a,a)}. \tag{5}$$

As a consequence, $(1-\gamma)V$ must be greater than 1. We shall call

$$\mathcal{D} := \{a \in \mathbb{R} \mid \text{condition (5) is fulfilled}\} \tag{6}$$

the *feasible trait domain* and assume that all traits (unless specified otherwise) are in $\mathcal{D}$.

We assume the system to be at this equilibrium when the mutant with trait $A \in \mathcal{D}$ arises. Thus, we consider the following initial conditions

$$(x(t=0), y(t=0), z(t=0), u(t=0)) = (\bar{x}(a), \epsilon_y, \bar{z}(a), \epsilon_u)$$

where $\epsilon_y, \epsilon_u > 0$ are infinitely small. We then place ourselves at $t = 0^+$. At first order, $x(t)$ and $z(t)$ are constant, therefore the dynamics of the invading population are driven by the linear system

$$\begin{cases} \frac{dy}{dt}\Big|_{t=0^+} &= (b(A) - d(A) - c(a,A)\bar{x}(a) - \eta(A)\bar{z}(a))\epsilon_y + \eta(a)\psi(a,A)\bar{x}(a)\epsilon_u \\ \\ \frac{du}{dt}\Big|_{t=0^+} &= \gamma V \eta(A)\bar{z}(a)\epsilon_y - (d_v + \eta(a)\bar{x}(a))\epsilon_u. \end{cases} \tag{7}$$

Invasion success is determined by the eigenvalues of this linear system. If both values are real and negative, the resident equilibrium is stable, and invasion fails. On the contrary, if both values are real (being conjugated) and one is positive, invasion succeeds. The eigenvalues, denoted by $\lambda_+$ and $\lambda_-$, are given by

$$\lambda_\pm = \frac{1}{2}\left(S(A,a) - (d_v + \eta(a)\bar{x}(a)) \pm \sqrt{\Delta}\right) \tag{8}$$

where

$$S(A,a) = r(A) - c(a,A)\bar{x}(a) - \eta(A)\bar{z}(a) \tag{9}$$

and

$$\Delta = (S(A,a) + (d_v + \eta(a)\bar{x}(a)))^2 + 4(\gamma\eta(a)\eta(A)\psi(a,A)V\bar{x}(a)\bar{z}(a)).$$

Here, $S(A, a)$ represents the invasion fitness in the absence of transduction, meaning when $\psi(a, A) = 0$. As expected intuitively, invasion fitness is higher if the mutant intrinsic growth rate is larger, or the competition or infection pressure experienced by the mutant from the resident population is weaker.

Since both additive terms in $\Delta$ are positive, we have $\Delta \geq 0$ (i.e. all eigenvalues are real). Both eigenvalues being real, we can focus on the greater of the two, which is $\lambda_+$. Then there are two cases: either $\lambda_+$ is negative, implying that the resident equilibrium is stable, or it is positive, implying that the resident equilibrium is unstable and invasion succeeds.

If $\Delta > (S(A, a) - (d_v + \eta(a)\bar{x}(a)))^2$, then $\lambda_+ > 0$. By rewriting $\Delta$ as

$$\Delta = (S(A, a) - (d_v + \eta(a)\bar{x}(a)))^2 + 4((d_v + \eta(a)\bar{x}(a))S(A, a) + \gamma\eta(a)\eta(A)\psi(a, A)V\bar{x}(a)\bar{z}(a)).$$

we can conclude that $\lambda_+ > 0$ if

$$S(A, a) + \frac{\gamma}{1-\gamma} \times \eta(A)\bar{z}(a) \times \psi(a, A) > 0. \tag{10}$$

Reciprocally, if $\lambda_+ > 0$, either $S(A, a) - (d_v + \eta(a)\bar{x}(a)) > 0$, then in particular $S(A, a) > 0$ and (10) is satisfied; or $-\sqrt{\Delta} < S(A, a) - (d_v + \eta(a)\bar{x}(a)) < 0$, and after some algebra we can conclude that (10) is also satisfied.

We then obtain invasion fitness with transduction, denoted by $S_T(A, a)$:

$$S_T(A, a) = S(A, a) + \frac{\gamma}{1-\gamma}\eta(A)\bar{z}(a)\psi(a, A). \tag{11}$$

If $S_T(A, a) > 0$, a mutant individual with trait $A$ can invade a resident population with trait $a$, otherwise invasion does not occur. From this two results follow:

- A mutant's invasion fitness can be negative in the absence of transduction ($S(A, a) < 0$) and yet positive with transduction ($S_T(A, a) > 0$). In other words, a mutation selected against in the absence of transduction may still be favored when transduction operates.
- For any trait value $a \in \mathbb{R}$ we have

$$S_T(a, a) = \frac{\gamma}{1-\gamma}\eta(a)\bar{z}(a)\psi(a, a) > 0. \tag{12}$$

More generally, individual cells may have different genotypes, $\mathcal{M}_i \neq \mathcal{M}_j$, and yet identical phenotypes, $a_i = a_j$. When this is the case, transduction will create a new lineage with potentially a different value for $a_{i\cup j}$, unless one population is a direct descendant of the other.

#### Back invasion

Given the conditions under which a mutant population can invade a resident population, we now ask how the resident population fares when rare in the mutant population, once the latter has spread. This case is useful to determine whether invasion by the mutant may result in a coexistence equilibrium between mutant and resident, or whether we can work under an *invasion implies fixation* principle.

The same calculations as before lead to the following expression for the back invasion fitness $\tilde{S}_T$:

$$\tilde{S}_T(a, A) = S(a, A) - \frac{\gamma}{1-\gamma}\eta(a)\bar{z}(A)\psi(a, A). \tag{13}$$

Values of $a, A$ may exist for which both $S_T$ and $\tilde{S}_T$ would be positive. But if we work under the assumption that mutations steps are small, i.e. that $A = a + \epsilon$ with $\epsilon \ll 1$, and that the population is distant from the evolutionary singluarity, then we can expound the invasion fitnesses as follows:

$$\begin{cases} S_T(A, a) = & \frac{\gamma}{1-\gamma}\eta(a)\bar{z}(a)\psi(a, a) + o(\epsilon) & > 0 \\ \tilde{S}_T(a, A) = & -\frac{\gamma}{1-\gamma}\eta(a)\bar{z}(a)\psi(a, a) + o(\epsilon) & < 0. \end{cases}$$

Based on Geritz et al. (2002) and Geritz (2005), we conclude that for small mutations, no coexistence is possible between mutant and resident populations, and that a (small) mutation is always at a selective advantage due to transduction. Introducing transduction alters the mathematical framework of Geritz et al. 2002 and Geritz 2005, but only so slightly, and numerical evidence supports the absence of coexistence. For $\psi_0 = 0$ (i.e. no transduction), the theoretical framework holds true and the results can be directly applied.

Under the small mutation assumption, we prove in the same way as we did for the resident equilibrium that the new equilibrium (since $A \in \mathcal{D}$) is given by

$$
\begin{cases}
\bar{y}(A) &= \frac{d_v}{\eta(A)((1-\gamma)V-1)} \\
\bar{z}(A) &= \frac{1}{\eta(A)}\left(r(A) - \frac{d_v}{\eta(A)((1-\gamma)V-1)}c(A,A)\right) \\
\bar{u}(A) &= \frac{\gamma}{1-\gamma}\bar{z}(A).
\end{cases}
\tag{14}
$$

### Supplementary note S3: Probability and characteristic time of invasion for a two-type population of bacteria with viruses

In the previous section, we found the condition for a mutant genotype of trait $A$ to invade a resident population with trait $a$. However, due to the stochastic nature of the process, even when a mutant genotype satisfies equation (10), invasion is not guaranteed. In this section, we derive the probability of invasion of a mutant genotype and the characteristic time it takes for an invasion to unfold. The derivation is largely based on Champagnat, Ferrière, et al. (2006).

#### Coupling of the process with a simple branching process

The system is set at the resident equilibrium (4), denoted by $(\bar{x}(a), 0, \bar{z}(a), 0)$. We assume that the mutation is small enough so that no coexistence between mutant and resident populations is possible. To the resident equilibrium we apply a small perturbation $(Y, U)$ assumed to be negligible compared to the other values of the equilibrium. For $K$ large enough and some population size threshold $\epsilon > 0$, we divide an invasion event into three distinct phases, as in Champagnat (2006):

1. The mutant population $Y^K$ either reaches the threshold size $\epsilon$ or goes extinct.
2. If the mutant population is not extinct, a new equilibrium will be reached following the deterministic approximation until the resident population reaches the threshold size $\epsilon$.
3. The resident population goes extinct[2].

The idea here is the following: we want to prove that when $K \to +\infty$, the process describing the invading population $(Y^K, U^K)$ tends to a branching process until the total population reaches a threshold size $\epsilon > 0$, while $X^K, Z^K$ stay relatively constant at their dynamical equilibrium.

[2]This occurs because the small mutation assumption implies no coexistence between mutant and resident

#### The ecological equilibrium of the resident population

We have seen earlier that for $K$ going to infinity, the population process is deterministic, given by (2). This implies, as we show here, that the resident bacterial population and the viral population stay at ecological equilibrium during the first phase of mutant invasion.

First, we know that during the first phase, the mutant population $Y^K$ has not reached the threshold $\epsilon$ yet. If we refer to the dynamical equation, this means that, at first order with respect to $\epsilon$, the resident bacterial and viral populations stay at ecological equilibrium $(\bar{x}(a), \bar{z}(a))$. We also know that the jumps of the stochastic process are of order $O(1/K)$. Since $K$ is large and the ecological equilibrium $(\bar{x}(a), \bar{z}(a))$ is term-by-term nonzero, we can conclude that at order $O(1/K)$, resident bacterial and virion populations stay at ecological equilibrium $(\bar{x}(a), \bar{z}(a))$. Therefore, we can assume that during the first phase, the resident bacterial and viral populations remain constant in size at ecological equilibrium.

#### Trajectorial representation of the invading population during the first phase

We define the following Poisson point measures:

- **Birth of a mutant:** $\mathcal{N}_1(ds, du)$ on $\mathbb{R}_+ \times \mathbb{R}_+$ with intensity measure $ds\ du$
- **Death of a mutant:** $\mathcal{N}_2(ds, du)$ on $\mathbb{R}_+ \times \mathbb{R}_+$ with intensity measure $ds\ du$
- **Infection of a mutant:** $\mathcal{N}_3(ds, du)$ on $\mathbb{R}_+ \times \mathbb{R}_+$ with intensity measure $ds\ du$
- **Transduction of a resident:** $\mathcal{N}_4(ds, du)$ on $\mathbb{R}_+ \times \mathbb{R}_+$ with intensity measure $ds\ du$
- **Death of a GTP:** $\mathcal{N}_5(ds, du)$ on $\mathbb{R}_+ \times \mathbb{R}_+$ with intensity measure $ds\ du$

Then we describe the mutant process through the following trajectorial representation

$$\begin{pmatrix} Y_t^K \\ U_t^K \end{pmatrix} = \begin{pmatrix} Y_0^K \\ U_0^K \end{pmatrix} + \int_0^t \int_{\mathbb{R}} \begin{pmatrix} +1/K \\ 0 \end{pmatrix} \mathbb{1}_{u \leq Y_{s-}^K b(A)} \mathcal{N}_1(ds, du)$$

$$+ \int_0^t \int_{\mathbb{R}} \begin{pmatrix} -1/K \\ 0 \end{pmatrix} \mathbb{1}_{u \leq Y_{s-}^K (d(A)+c(a,A)\bar{x}(a)+c(A,A)Y_{s-}^K)} \mathcal{N}_2(ds, du)$$

$$+\int_0^t\int_{\mathbb{R}}\sum_{i=0}^{V}\begin{pmatrix}-1/K\\ i/K\end{pmatrix}\mathbb{1}_{Y^K_{s-}\sum_{j=0}^{i-1}p_i^{(V,\gamma)}\eta(A)\bar{z}(a)\ <u\ \leq\ Y^K_{s-}\sum_{j=0}^{i}p_i^{(V,\gamma)}\eta(A)\bar{z}(a)}\ \mathcal{N}_3(ds,du)$$

$$+\int_0^t\int_{\mathbb{R}}\begin{pmatrix}+1/K\\ -1/K\end{pmatrix}\mathbb{1}_{u\ \leq\ U^K_{s-}\ \ \eta(A)\bar{z}(a)}\ \mathcal{N}_4(ds,du)$$

$$+\int_0^t\int_{\mathbb{R}}\begin{pmatrix}0\\ -1/K\end{pmatrix}\mathbb{1}_{u\ \leq\ U^K_{s-}\ \ (d_v+\eta(A)Y^K_{s-})}\ \mathcal{N}_5(ds,du).$$

#### Bounding our process with branching processes

We now show that there exist two branching processes $(\widetilde{Y}^{K,1},\widetilde{U}^{K,1}),(\widetilde{Y}^{K,2},\widetilde{U}^{K,2})$ such that

$$\forall t\in\mathbb{R},\quad \begin{aligned}\widetilde{Y}^{K,1}_t\leq Y^K_t\leq \widetilde{Y}^{K,2}_t\\ \widetilde{U}^{K,1}_t\leq U^K_t\leq \widetilde{U}^{K,2}_t.\end{aligned}\tag{15}$$

Using the same Poisson point measures and the same initial condition as in the previous section, we define the following processes

$$\begin{pmatrix}\widetilde{Y}^{K,1}_t\\ \widetilde{U}^{K,1}_t\end{pmatrix}=\begin{pmatrix}Y^K_0\\ U^K_0\end{pmatrix}+\int_0^t\int_{\mathbb{R}}\begin{pmatrix}+1/K\\ 0\end{pmatrix}\mathbb{1}_{u\ \leq\ \widetilde{Y}^{K,1}_{s-}\ \ b(A)}\ \mathcal{N}_1(ds,du)$$

$$+\int_0^t\int_{\mathbb{R}}\begin{pmatrix}-1/K\\ 0\end{pmatrix}\mathbb{1}_{u\ \leq\ \widetilde{Y}^{K,1}_{s-}\ \ (d(A)+c(a,A)\bar{x}(a)+c(A,A)\epsilon)}\ \mathcal{N}_2(ds,du)$$

$$+\int_0^t\int_{\mathbb{R}}\sum_{i=0}^{V}\begin{pmatrix}-1/K\\ i/K\end{pmatrix}\mathbb{1}_{\widetilde{Y}^{K,1}_{s-}\sum_{j=0}^{i-1}p_i^{(V,\gamma)}\eta(A)\bar{z}(a)\ <u\ \leq\ \widetilde{Y}^{K,1}_{s-}\sum_{j=0}^{i}p_i^{(V,\gamma)}\eta(A)\bar{z}(a)}\ \mathcal{N}_3(ds,du)$$

$$+\int_0^t\int_{\mathbb{R}}\begin{pmatrix}+1/K\\ -1/K\end{pmatrix}\mathbb{1}_{u\ \leq\ \widetilde{U}^{K,1}_{s-}\ \ \eta(A)\bar{z}(a)}\ \mathcal{N}_4(ds,du)$$

$$+\int_0^t\int_{\mathbb{R}}\begin{pmatrix}0\\ -1/K\end{pmatrix}\mathbb{1}_{u\ \leq\ \widetilde{U}^{K,1}_{s-}\ \ (d_v+\eta(A)\epsilon)}\ \mathcal{N}_5(ds,du)$$

and

$$\begin{pmatrix} \widetilde{Y}_t^{K,2} \\ \widetilde{U}_t^{K,2} \end{pmatrix} = \begin{pmatrix} Y_0^K \\ U_0^K \end{pmatrix} + \int_0^t \int_{\mathbb{R}} \begin{pmatrix} +1/K \\ 0 \end{pmatrix} \mathbb{1}_{u \,\leq\, \widetilde{Y}_{s^-}^{K,2}\, b(A)}\, \mathcal{N}_1(ds, du)$$

$$+ \int_0^t \int_{\mathbb{R}} \begin{pmatrix} -1/K \\ 0 \end{pmatrix} \mathbb{1}_{u \,\leq\, \widetilde{Y}_{s^-}^{K,2}\, (d(A)+c(a,A)\bar{x}(a))}\, \mathcal{N}_2(ds, du)$$

$$+ \int_0^t \int_{\mathbb{R}} \sum_{i=0}^{V} \begin{pmatrix} -1/K \\ i/K \end{pmatrix} \mathbb{1}_{\widetilde{Y}_{s^-}^{K,2} \sum_{j=0}^{i-1} p_i^{(V,\gamma)} \eta(A)\bar{z}(a) \,<u\, \leq\, \widetilde{Y}_{s^-}^{K,2} \sum_{j=0}^{i} p_i^{(V,\gamma)} \eta(A)\bar{z}(a)}\, \mathcal{N}_3(ds, du)$$

$$+ \int_0^t \int_{\mathbb{R}} \begin{pmatrix} +1/K \\ -1/K \end{pmatrix} \mathbb{1}_{u \,\leq\, \widetilde{U}_{s^-}^{K,2}\, \eta(A)\bar{z}(a)}\, \mathcal{N}_4(ds, du)$$

$$+ \int_0^t \int_{\mathbb{R}} \begin{pmatrix} 0 \\ -1/K \end{pmatrix} \mathbb{1}_{u \,\leq\, \widetilde{U}_{s^-}^{K,2}\, d_v}\, \mathcal{N}_5(ds, du).$$

The only differences arise from representing the death of a mutant cell and the death of a GTP. The second order terms have been either replaced by $\epsilon$ (for the lower bound; indeed, by definition of the first phase, $Y_t^K \leq \epsilon$) or completely removed (for the upper bound).

We now check that the following is true:

$$\forall t \in \mathbb{R},\ \widetilde{Y}_t^{K,1} \leq Y_t^K \leq \widetilde{Y}_t^{K,2},$$
$$\widetilde{U}_t^{K,1} \leq U_t^K \leq \widetilde{U}_t^{K,2}.$$

*Proof.* We show that if at time $t_1 \in \mathbb{R}_+$, (15) is satisfied, then it will be satisfied for all times $t > t_1$. Let us call $T$ the first event that occurs after $t_1$. Since the Poisson point measures are the same, we can conclude that we will be studying the same point in one of the PPM $\mathcal{N}_i$ for $i \in 1, ..., 5$. The possible events are:

- **Birth of a mutant:** From the assumptions we have

$$\widetilde{Y}_{T^-}^{K,1} \leq Y_{T^-}^K \leq \widetilde{Y}_{T^-}^{K,2}$$

so

$$\widetilde{Y}_{T^-}^{K,1} b(A) \leq Y_{T^-}^K b(A) \leq \widetilde{Y}_{T^-}^{K,2} b(A)$$

which means that a jump of $+1/K$ for $\widetilde{Y}^{K,1}$ (i.e. $u \leq \widetilde{Y}^{K,1}b(A)$) would imply a jump for $Y^K$ which would also imply a jump for $\widetilde{Y}^{K,2}$. In all cases, the order is respected.

- **Death of a mutant:** Here, the previous argument does not apply. Indeed, let us focus on the relationship between $\widetilde{Y}^{K,1}_{T^-}$ and $Y^K_{T^-}$. We know that

$$\widetilde{Y}^{K,1}_{T^-} \leq Y^K_{T^-}$$

but

$$d(A) + c(a,A)\bar{x}(a) + c(A,A)\epsilon \geq d(A) + c(a,A)\bar{x}(a) + c(A,A)Y^K_{T^-} \tag{16}$$

so there could be instances where $Y^K$ would jump of $-1/K$ but not $\widetilde{Y}^{K,1}$. We then need to distinguish between two cases:

  - If $Y^K_{T^-} > \widetilde{Y}^{K,1}_{T^-}$ then we know that $Y^K_{T^-} - 1/K \geq \widetilde{Y}^{K,1}_{T^-}$, since $Y^K, \widetilde{Y}^{K,1} \in \mathbb{N}/K$, so the order is retained.
  - If $Y^K_{T^-} = \widetilde{Y}^{K,1}_{T^-}$, then since (16) we can infer

$$\widetilde{Y}^{K,1}_{T^-}(d(A) + c(a,A)\bar{x}(a) + c(A,A)\epsilon) \geq Y^K_{T^-}(d(A) + c(a,A)\bar{x}(a) + c(A,A)Y^K_{T^-})$$

    so a jump for $Y^K$ would imply a jump for $\widetilde{Y}^{K,1}$. Again the order is retained.

  The same arguments can be made between $Y^K$ and $\widetilde{Y}^{K,2}$, so we can conclude that the order is respected.

- **Other events:** The three other events that may occur can be treated in the same way as we just did for the first two cases.

Since we have the same initial conditions, we can conclude that (15) is verified for $t = 0$, meaning we have proved it for all $t > 0$. We thus have constructed two branching processes such that

$$\forall t \in \mathbb{R}, \ \widetilde{Y}^{K,1}_t \leq Y^K_t \leq \widetilde{Y}^{K,2}_t$$
$$\widetilde{U}^{K,1}_t \leq U^K_t \leq \widetilde{U}^{K,2}_t.$$

□

##### Conclusion

We just proved that for every $K \in \mathbb{N}^*$, every $\epsilon > 0$ and especially every $\omega$ ($\omega$ being the realization of the Poisson point measures) we have

$$\begin{aligned} \forall t \in \mathbb{R}, \widetilde{Y}_t^{K,1} \leq Y_t^K \leq \widetilde{Y}_t^{K,2} \\ \widetilde{U}_t^{K,1} \leq U_t^K \leq \widetilde{U}_t^{K,2} \end{aligned}$$

where $\forall t \in \mathbb{R}, \lim_{\epsilon \to 0} \widetilde{Y}_t^{K,1} = \widetilde{Y}_t^{K,2}$ (resp. for $\widetilde{U}^{K,i}$).

We can then conclude that, when $K$ is large and $\epsilon \to 0$, the process $(Y^K, U^K)_t$ converges with high probability towards the two-type branching process with the following transitions:

| **Next state** | **Rate** | **Event** |
|---|---|---|
| $(Y+1, U)$ | $b(A)Y$ | *Birth* |
| $(Y-1, U)$ | $(d(A) + c(a, A)\bar{x}(a))Y$ | *Death* |
| $(Y+1, U-1)$ | $\eta(a)\psi(a, A)\bar{x}(a)U$ | *Transduction* |
| $(Y-1, U+k)$ | $\eta(A)\bar{z}(a)p_k^{(V,\gamma)}Y$ | *Infection* |
| $(Y, U-1)$ | $d_v U + \eta(a)(1-\psi(a, A))\bar{x}(a)U$ | *Death* |

The same reasonning can be applied to the third phase with individuals of type $Y, Z, U$ being at ecological equilibrium and the population of type $X$ going to extinction. We then find a simple birth and death branching process which we will study in a subsequent section.

#### Probability of invasion

Once the process reaches the threshold size $\epsilon$, the large population limit allows us to conclude that the invasion is successful (since $S_T(A, a) > 0$). Therefore, the probability of invasion of the mutant population is the same as the probability that the coupled two-type branching process described above diverges towards infinity. To calculate this probability, we introduce

$u_{i,j} = \mathbb{P}((Y,U)$ will go extinct $|Y = i, U = j)$ and we use the notations $u_1 = u_{1,0}$ and $u_2 = u_{0,1}$. We have

$$\forall i, j \geq 0, u_{i,j} = (u_1)^i \times (u_2)^j. \tag{17}$$

When analysing the process, we find that $(u_1, u_2)$ satisfies

$$\begin{cases} u_1 & = q_1 u_1^2 + \sum_{k=0}^{V} q_2 p_k^{(V,\gamma)} (u_2)^k + (1 - q_1 - q_2) \\ \\ u_2 & = q_3 u_1 + (1 - q_3) \end{cases} \tag{18}$$

where

$$q_1 = \frac{b(A)}{b(A) + d(A) + c(a,A)\bar{x}(a) + \eta(A)\bar{z}(a)}$$

$$q_2 = \frac{\eta(A)\bar{z}(a)}{b(A) + d(A) + c(a,A)\bar{x}(a) + \eta(A)\bar{z}(a)}$$

$$q_3 = \frac{\eta(a)\psi(a,A)\bar{x}(a)}{\eta(a)\bar{x}(a) + d_v} = \frac{\psi(a,A)}{(1-\gamma)V}.$$

We recall that (5) implies $(1-\gamma)V > 1$, and therefore $q_3 < 1$.

Assuming that an invasion event starts off with one mutant cell, the probability of invasion is $p_T(A,a) = 1 - u_1$. The corresponding term in equation (18) (first line) is equal to

$$u_1 = q_1 u_1^2 + q_2 (1 - \gamma + \gamma u_2)^V + (1 - q_1 - q_2).$$

We note that $u_2 = 1 - q_3(1 - u_1) = 1 - q_3 p_T(A,a)$ and substitute $u_1$ with $1 - p_T(A,a)$. We then find that $p_T(A,a)$ satisfies the following equation

$$b(A)\ p_T(A,a)^2 - S(A,a)\ p_T(A,a) + \eta(A)\ \bar{z}(a) \left(1 - \frac{\gamma\psi(a,A)}{(1-\gamma)V}\ p_T(A,a)\right)^V - \eta(A)\ \bar{z}(a) = 0.$$

For the sake of legibility, we will continue to use $q_3$ for the remainder of the proof.

Interestingly, when transduction does not occur ($\psi(a,A) = 0$), we have $q_3 = 0$ and the result becomes

$$p(A,a) = \frac{[S(A,a)]_+}{b(A)}. \tag{19}$$

The result is similar to the case of a simple birth-death branching process. This is because the GTP population has no impact when $\psi(a,A) = 0$.

We now focus on proving the existence and uniqueness of a root $p \in (0;1)$ to the following polynomial, $Q$, under the condition $S_T(A,a) > 0$, where $S_T(A,a)$ is given by equation (11):

$$Q = b(A) \times X^2 - S(A,a) \times X + \eta(A)\bar{z}(a)(1 - \gamma q_3 \times X)^V - \eta(A)\bar{z}(a).$$

The claim follows from *Descartes' Rule of Signs* (Descartes 1637) (if the terms of a single-variable polynomial with real coefficients are ordered by descending variable exponent, then the number of positive roots of the polynomial is either equal to the number of sign differences between consecutive nonzero coefficients, or is less than it by an even number). To apply Descartes' Rule, we change variables:

$$\xi = 1 - \gamma\ q_3\ X$$

Since $\gamma < 1, q_3 < 1$, we have $X \in [0;1]$ if and only if $\xi \in [1 - \gamma q_3; 1] \subset [0;1]$. Polynomial $Q$ becomes $\tilde{Q}$ given by

$$\tilde{Q} = \eta(A)\bar{z}(a)\xi^V + \frac{b(A)}{\gamma^2 q_3^2}\xi^2 + \frac{1}{\gamma q_3}\left(S(A,a) - \frac{2b(A)}{\gamma q_3}\right)\xi - \eta(A)\bar{z}(a) - \frac{S(A,a)}{\gamma q_3} + \frac{b(A)}{\gamma^2 q_3^2}.$$

We wrote $\tilde{Q}$ terms in the order of decreasing exponents. The first two coefficients are non-negative. Since the polynomial has only four coefficients, there can be at most two sign switches between consecutive coefficients (between the second and third, and between the third and fourth). Descartes' Rule signs then assures us that $\tilde{Q}$ will have **at most** two positive roots.

Now we show that we can find **at least** two roots of $\tilde{Q}$ in $[1 - \gamma q_3; 1]$ under the condition $S(A,a) > 0$. From this we will conclude that there are exactly two positive roots of $\tilde{Q}$, and that they are both in $[1 - \gamma q_3; 1]$. One can easily check the following:

$$\tilde{Q}(1) = 0$$

$$\tilde{Q}(1-\gamma q_3) = d(A) + c(a,A)\bar{x}(a) + \eta(A)\ \bar{z}(a)(1-\gamma q_3)^V > 0$$

$$\tilde{Q}'(1) = S(A,a) + \eta(A)\bar{z}(a)V\gamma q_3 = S(A,a) > 0$$

Therefore, there is a root of $\tilde{Q}$ in $(1-\gamma q_3; 1)$. Since 1 is also a root, we did find at least two roots of $\tilde{Q}$ in $[1-\gamma q_3; 1]$. This implies that we have exactly two positive roots, both in $[1-\gamma q_3; 1]$. Polynomial $Q$ has exactly two roots in $[0; 1]$ when $S_T(A,a) > 0$, one being zero and the other one being $p_T(A,a)$.

We now compare this root to $p(A,a)$, defined in equation (19). If $S(A,a) \leq 0$, $p(A,a) = 0$ and then $p_T(A,a) \geq p(A,a)$. If $S(A,a) > 0$, then $p(A,a) = \frac{S(A,a)}{b(A)}$. Evaluating $Q$ at $p(A,a)$ yields

$$Q(p(A,a)) = \eta(A)\bar{z}(a)(1-\gamma q_3\ p(A,a))^V - \eta(A)\bar{z}(a) < 0.$$

$Q'(0) < 0$ implies $p_T(A,a) > p(A,a)$. In sum, we have shown that transduction always increases the probability of invasion of a mutant.

#### A useful approximation

An analytical approximation for the probability of invasion can be found in the case $V \gg 1$, which would lead to this interpretation of the defining equat. In this case, $p_T(A,a)$ is the solution to

$$b(A)\ p_T(A,a)^2 - S(A,a)\ p_T(A,a) + \eta(A)\bar{z}(a)\ \exp\left(-\frac{\gamma}{1-\gamma}\psi(a,A)p_T(A,a)\right) - \eta(A)\bar{z}(a) = 0.$$

Then if we assume that $\gamma\psi(a,A)p_T(A,a) \ll 1$ (which appears to be the case for values as high as 0.1 for two or three of the three factors), we can further approximate the equation:

$$b(A)\ p_T(A,a)^2 - \underbrace{\left(S(A,a) + \frac{\gamma}{1-\gamma}\psi(a,A)\eta(A)\bar{z}(a)\right)}_{=S_T(A,a)} p(A,a) = 0.$$

This leads us to the approximated expression $\widetilde{p}_T(A,a)$ for the probability of invasion:

$$\widetilde{p}_T(A,a) = \frac{[S_T(A,a)]_+}{b(A)}. \tag{20}$$

We note that transduction does not fundamentally change the mathematical expression of the probability of invasion (invasion fitness divided by the mutant birth rate). A similar expression was found also in the case of genetic mobility due to conjugation (Billiard et al. 2016).

##### Invasion probability of a neutral mutation

Since $S_T(a,a) > 0$, we have $p_T(a,a) > 0$, i.e. a positive invasion probability for a mutant that is phenotypically (but not genetically) identical to the resident genotype.

#### Characteristic time of invasion

As in the previous section, we divide the invasion process in the same three distinct phases:

1. The time it takes for the mutant population to reach the threshold size $\epsilon K$ can be calculated as the time that a two-type linear birth and death process takes for its first component to reach the same threshold.

2. The deterministic phase takes a constant time, thus of order $O(1)$.

3. The time it takes for the resident population to go extinct once it reaches threshold size $\epsilon K$ can be calculated as the time of extinction for a single type linear birth and death process.

The calculations are similar to what was done in previous studies (Billiard et al. 2016, Méléard 2016). We conclude that the characteristic time of invasion $T_{inv}$ is of order $O(\log(K))$, and that it almost surely does not diverge from this order of magnitude.

### Supplementary note S4: Trait Substitution Sequence and canonical equation

This section is developed under the *Invasion Implies Fixation* (IFF) set of assumptions. This means that every invasion event may only have one of two outcomes: either the mutant replaces the resident population, or it goes extinct. This has not been rigorously proved in our case, but there is no hint from numerical simulations running against this assumption.

We recall that the phenotype space is defined as the *valid trait domain*

$$\mathcal{D} := \{a \in \mathbb{R} \mid \text{condition (5) is fulfilled}\}. \tag{21}$$

We denote the mutation rate of the bacterial population by $\mu_K$. The mathematical derivation assumes rare mutations, meaning $\mu_K \to 0$ when $K \to +\infty$. We make the further assumption that

$$\forall \gamma > 0,\ \log K \ll \frac{1}{K\mu_K} \ll \exp(\gamma K) \tag{22}$$

and we rescale time according to

$$t' := \frac{t}{K\mu_K}.$$

To justify the first condition in equation (22), we recall that the characteristic time of invasion is of order $O(\log K)$ and never exceeds this order of magnitude (see previous section). This implies that on the timescale set by $t'$, invasion events resolve instantaneously, and two mutations occurring at thr same instant is highly improbable. Under the *Invasion Implies Fixation* principle, each invasion event produces a "winner" and a "loser", and thus the model describes the population adaptation dynamics on the $t'$ timescale as if only one trait value was present in the population at any time.

The second condition in equation (22) is supported by large deviation theory (Freidlin and Wentzell 1998, Feng and Kurtz 2006, Dupuis and Ellis 1997). We define $\Omega(a) = (\bar{x}(a), \bar{z}(a))$ the equilibrium point, $\mathcal{O} \in \mathbb{R}^2$ an open set which contains $\Omega(a)$ and $w_t^K = (X_t^K, Z_t^K)$, and

$$T^K = \inf\left\{t \geq 0; w_t^K \notin \mathcal{O}\right\}.$$

Then there exists some value $\gamma > 0$ such that, for any compact subset $C$ of $\mathcal{O}$, we have

$$\lim_{K\to+\infty} \sup_{z\in C} \mathbb{P}_z^K(T^K < e^{\gamma K}) = 0. \tag{23}$$

Thus, considering a process unfolding between these two timescales, i.e. on the $t' := t/K\mu_K$ timescale, we conclude that events involving the occurrence of two or more mutations at the same time are very rare, and that population sizes will stay at ecological equilibrium at any given time. We can then consider invasion events as instantaneous, occurring at the population rate of mutation, with a probability of success given by $p_T$ (see previous section).

On this evolutionary timescale, the trait dynamics is a jump process, called the *Trait Substitution Sequence* (TSS). Jumps from trait $a \in \mathcal{D}$ to trait $A \in \mathcal{D}$ occur at rate

$$p_T(A, a)\ \bar{x}(a)b(a)m(a, dA) \tag{24}$$

where $\bar{x}(a)$ is the equilibrium point of the resident population of trait $a$; $m(a, dA)$ is the mutation kernel from trait $a$ to trait $A$ that we can write

$$m(a, dA) = m(A - a)\ dA.$$

On the evolutionary timescale, the population birth rate, $\bar{x}(a)b(a)$, measures the probability of a mutant birth per unit time in a homogeneous population of trait $a$. We can then write the jump rate from $a$ to $A$ as

$$\chi(a, A) := p_T(A, a)\ \bar{x}(a)b(a)m(A - a). \tag{25}$$

From our assumptions it follows that there exists $C_\chi \in \mathbb{R}$ such as $\forall a, A \in \mathcal{D}, \chi(a, A) \leq C_\chi$.

#### Convergence to the canonical equation

We now focus on small mutation steps, the difference $A - a$ being of order $\epsilon > 0$. Under this assumption we have

$$\int_{\mathcal{D}} g(A)m_\epsilon(a, dA) = \int_{\mathcal{D}} g(a + \epsilon h)m(a, dh) = \int_{\mathcal{D}} g(a + \epsilon h)m(a, h)dh.$$

To capture the adaptation dynamics on a longer timescale, we take the limit $\epsilon \to 0$ and rescale time. The order in which the double rescaling is done might matter. Here we show that different outcomes may ensue, depending on the shape of the mutation kernel.

**Symmetrical mutation kernel**

In this case, $\forall a \in \mathcal{D}, h \mapsto m(a,h)$ is assumed symmetrical with respect to $h$. As we shall see, the following time rescaling is relevant:

$$\xi_t^\epsilon = a_{t/\epsilon^2}^\epsilon.$$

The rescaling in $\epsilon^2$ is a natural one in the analysis of the TSS, but is usually combined with the fact that $\forall a \in \mathcal{D}, p(a,a) = 0$. Here, a meaningful $\epsilon^2$ rescaling is obtained provided the mutation kernel is symmetrical.

Let $T \in \mathbb{R}_+$ be some large time horizon. We restrict the analysis of the process to the interval $[0;T]$, meaning that $\xi^\epsilon \in \mathbb{D}([0,T],\mathbb{R}_+)$. The $\epsilon^2$ rescaling together with the technical assumption $\mathbb{E}\left[(\xi_0^\epsilon)^3\right] < +\infty$ imply that the TSS converges in law to the following stochastic equation

$$d\xi_t = \bar{b}(\xi_t)\ dt + \sigma(\xi_t)\ dB_t \tag{26}$$

with

$$\bar{b}(\xi_t) = \bar{x}(\xi_t)b(\xi_t)\sigma_0^2(\xi_t)\partial_1 p_T(\xi_t,\xi_t) \tag{27}$$

$$\sigma(\xi_t) = \sigma_0(\xi_t)\sqrt{\bar{x}(\xi_t)b(\xi_t)p_T(\xi_t,\xi_t)} \tag{28}$$

where $\sigma_0(a)$ is the standard deviation of the mutation kernel when the resident trait is equal to $a$.

We make the further assumptions that the parameters of our model are regular enough to ensure that $\bar{b}$ and $\sigma$ are both Lipschitz on $\mathcal{D}$, the domain on which our analysis is valid.

The difference with classical stochastic equations comes from the stochastic term. This term only exists because $p_T(a,a) > 0$ for $a \in \mathcal{D}$, highlighting the importance of an explicit model of the genotype-phenotype map.

*Proof.* We have

$$a_t^\epsilon = a_0^\epsilon + \int_0^t \int_{\mathcal{D}} \int_{\mathbb{R}_+} (\alpha - a_{s^-}^\epsilon) \mathbb{1}_{u \leq \bar{x}(a_{s^-}^\epsilon) b(a_{s^-}^\epsilon) m(a_{s^-}^\epsilon, h) p_T(a_{s^-}^\epsilon + h\epsilon, a_{s^-}^\epsilon)} \mathcal{N}(du, dh, ds)$$

which can be re-written as

$$a_t^\epsilon = a_0^\epsilon + \epsilon \int_0^t \int_{\mathcal{D}} h\bar{x}(a_s^\epsilon) b(a_s^\epsilon) m(a_s^\epsilon, h) p_T(a_s^\epsilon + \epsilon h, a_s^\epsilon) dh ds + M_t^\epsilon$$

where

$$< M^\epsilon >_t = \epsilon^2 \int_0^t \int_{\mathcal{D}} h^2 \bar{x}(a_s) b(a_s) m(a_s^\epsilon, h) p_T(a_s^\epsilon + \epsilon h, a_s^\epsilon) dh ds.$$

By expanding $p$ to first order we obtain

$$\begin{aligned}
a_t^\epsilon &= a_0^\epsilon + \epsilon \int_0^t \int_{\mathcal{D}} h\bar{x}(a_s^\epsilon) b(a_s^\epsilon) m(h) \left( p_T(a_s^\epsilon, a_s^\epsilon) + \epsilon h \partial_1 p_T(a_s^\epsilon, a_s^\epsilon) + o(\epsilon) \right) dh ds + M_t^\epsilon \\
&= a_0^\epsilon + \epsilon \int_0^t \bar{x}(a_s^\epsilon) b(a_s^\epsilon) p_T(a_s^\epsilon, a_s^\epsilon) \underbrace{\int_{\mathcal{D}} h m(a_s^\epsilon, h) dh}_{=0 \text{ since } M \text{ is symmetrical}} ds \\
&+ \epsilon^2 \int_0^t \bar{x}(a_s^\epsilon) b(a_s^\epsilon) \partial_1 p_T(a_s^\epsilon, a_s^\epsilon) \int_{\mathcal{D}} h^2 m(a_s^\epsilon, h) dh ds + o(\epsilon^2) + M_t^\epsilon \\
&= a_0^\epsilon + \epsilon^2 \int_0^t \bar{x}(a_s^\epsilon) b(a_s^\epsilon) \partial_1 p_T(a_s^\epsilon, a_s^\epsilon) \sigma_0^2(a_s^\epsilon) ds + o(\epsilon^2) + M_t^\epsilon
\end{aligned}$$

with, for the martingale part:

$$< M^\epsilon >_t = \epsilon^2 \int_0^t \int_{\mathcal{D}} h^2 \bar{x}(a_s) b(a_s) m(a_s^\epsilon, h) p_T(a_s^\epsilon, a_s^\epsilon) dh ds + o(\epsilon^2).$$

This leads us to rescale time according to

$$\xi_t^\epsilon = a_{t/\epsilon^2}^\epsilon, \tag{29}$$

hence

$$\xi_t^\epsilon = \xi_0^\epsilon + \int_0^t \bar{x}(\xi_{s'}^\epsilon) b(\xi_{s'}^\epsilon) \partial_1 p_T(\xi_{s'}^\epsilon, \xi_{s'}^\epsilon) \sigma_0^2(\xi_s^\epsilon) ds' + \tilde{M}_t^\epsilon + o(1)$$

with

$$< \tilde{M}^\epsilon >_t = \int_0^t \bar{x}(\xi^\epsilon_{s'}) b(\xi^\epsilon_{s'}) p_T(\xi^\epsilon_{s'}, \xi^\epsilon_{s'}) \sigma_0^2(\xi^\epsilon_s) ds' + o(1). \tag{30}$$

When $\epsilon \to 0$, we have $< \tilde{M}^\epsilon > \to \int_0^t \sigma^2(\xi_s) ds$ which is non-zero, from where the stochastic part of the canonical equation follows. As we stated before, this special feature stems from the fact that $\forall a \in \mathcal{D}, p_T(a,a) > 0$. Other than this specificity, the analysis is similar to the case treated in Champagnat and Méléard 2011 and we refer to this work for a comprehensive proof of the convergence. □

##### Asymmetrical mutation kernel

Here the expectation of the mutational effect is

$$\mu_0(a) := \int_{\mathcal{D}} h m(a,h) dh$$

which is nonzero by assumption. By taking the same steps as before, we find

$$a^\epsilon_t = a^\epsilon_0 + \epsilon \int_0^t \bar{x}(a^\epsilon_s) b(a^\epsilon_s) p_T(a^\epsilon_s, a^\epsilon_s) \mu_0(a^\epsilon_s) ds + o(\epsilon) + M^\epsilon_t$$

with

$$< M^\epsilon >_t = \epsilon^2 \int_0^t \int_{\mathcal{D}} h^2 \bar{x}(a_s) b(a_s) m(a^\epsilon_s, h) p_T(a^\epsilon_s, a^\epsilon_s) dh ds + o(\epsilon^2).$$

Here the natural rescaling is

$$\widetilde{\xi}^\epsilon_t = a^\epsilon_{t/\epsilon}. \tag{31}$$

In contrast to the symmetrical mutation case, this rescaling smoothes out the stochastic part of the adaptation process, which is of order $\epsilon^2$. The limit then is the solution to the following ordinary equation

$$\widetilde{\xi}_t = \widetilde{\xi}_0 + \int_0^t \bar{x}(\widetilde{\xi}_s) b(\widetilde{\xi}_s) p_T(\widetilde{\xi}_s, \widetilde{\xi}_s) \mu_0(\widetilde{\xi}_s) ds. \tag{32}$$

In this case, the long-term adaptation dynamics are driven deterministically by selection.

### Boundaries of the stochastic canonical equation

Here we assume that the mutation kernel is symmetrical, and thus the canonical equation is stochastic. We define $\mathcal{I} = (a_{\text{MIN}}; a_{\text{MAX}}) \in \mathcal{D}$ as the largest interval within $\mathcal{D}$ which contains $a_0$, the trait value at which the canonical equation is initialized. We have $-\infty \leq a_{\text{MIN}} < a_{\text{MAX}} \leq +\infty$, and for $z \in \mathcal{I}, \sigma(z) \neq 0$.

Let us define the explosion time $e = \lim_{n\uparrow\infty} \tau_n$ where $\tau_n = \inf\{t; \xi_t \notin [a_n; b_n]\}$ ($a_n, b_n$ being chosen such that $a_{\text{MIN}} < a_n < b_n < a_{\text{MAX}}$ and $a_n \downarrow a_{\text{MIN}}, b_n \uparrow a_{\text{MAX}}$). In other words, $e$ is the time it takes for the process to exit the acceptable domain for the first time. In this model, such an event would mean that the virus population has gone extinct, as $\bar{z}(a) = 0$ at the limits of $\mathcal{D}$.

Given $c \in \mathcal{I}$ we set

$$s(x) := \int_c^x \exp\left|-\int_c^y \frac{2\bar{b}(z)}{\sigma^2(z)} dz\right| dy \tag{33}$$

By applying Theorem 3.1 of Chapter VI in Ikeda and Watanabe (2014) we know that if

$$\lim_{y\downarrow a_{\text{MIN}}} s(y) = -\infty \quad \text{and} \quad \lim_{y\uparrow a_{\text{MAX}}} s(y) = +\infty.$$

then

$$\mathbb{P}_{a_0}(e = \infty) = 1 \tag{34}$$

for every $a_0$. Since $p_T(z, z) \neq 0$ for $z \in \mathcal{D}$, we have

$$s(x) = \int_c^x \exp\left|-\int_c^y \frac{2\partial_1 p_T(z, z)}{p_T(z, z)} dz\right| dy.$$

Studying the convergence or divergence of such an integral is out of reach algebraically. But further numerical studies presented in the next section strongly suggest that under sufficient regularity conditions on $\bar{b}$ and $\sigma$, we do indeed find that $\lim_{y\downarrow a_{\text{MIN}}} s(y) = -\infty$ and $\lim_{y\uparrow a_{\text{MAX}}} s(y) = +\infty$. We can then assume $\mathbb{P}(e = \infty) = 1$. This means that with probability 1, the trait will not exit the valid trait domain $\mathcal{D}$, implying that there is no accidental extinction of the virus population.

### Supplementary note S5: Defining the stochasticity index $I$

In order to quantify how strong the stochastic perturbations are near the evolutionary singularity, we define a stochasticity index, $I$. Since our ultimate goal is to quantify the speed of convergence of the adaptation process with transduction, we need to provide support for the convergence of the process. If the process were dominated by transduction-induced stochasticity, the adaptive trajectories would not remain in a neighborhood of the evolutionary equilibrium.

First we quantitatively define a 'neighborhood' of the evolutionary singularity, $a^*$, in trait space. By construction, the trait value $a = a^*$ maximizes resource acquisition; in other words, this trait value maximizes the population carrying capacity $K(a)$, which is modeled as a gaussian curve with $K(a^*) = K_0$ (see **Supplementary Table 1**). We then define our reference neighborhood of $a^*$ as the range of $a$ values such that $K(a) > 0.99\ K_0$.

Then we define two characteristic times of interest for any pair of parameters $(\eta_0, \psi_0)$:

1. The *characteristic time of adaptation without transduction*, defined as $1/\alpha$ where $\alpha$ is the parameter used to fit an exponential curve to the TSS (with $\psi_0 = 0$). This characteristic time represents the time it takes for the TSS to complete roughly 2/3 of the distance to the evolutionary singularity.

2. The *mean sojourn time* in the evolutionary singularity neighborhood, with transduction (given $\psi_0 > 0$).

Both times were computed numerically based on the simulated TSS. The adaptation stochasticity index, $I$, is then computed as the ratio between the characteristic time of adaptation without transduction and the mean sojourn time with transduction. If $I \ll 1$, the stochastic part of the TSS drives the process away from equilibrium faster than the deterministic part pulls it back; stochasticity dominates the adaptation dynamics. If $I \gg 1$, the deterministic component of the TSS pulls the process near the evolutionary singularity faster than the stochastic part upsets it.

### Supplementary Table 1: Default simulation parameters

| Parameter | Form used | Numerical values |
|---|---|---|
| Population size order | $K$ constant | $K = 200$ |
| Trait range | $]a_{\text{INF}}, a_{\text{SUP}}[$ | $a_{\text{INF}} = -1.5, a_{\text{SUP}} = 1.5$ |
| Birth rate | $b(a) = b_0$ constant | $b_0 = 4$ day$^{-1}$ |
| Death rate | $d(a) = d_0$ constant | $d_0 = 0.5$ day$^{-1}$ |
| Competition rate | $c(A, a) = \frac{\exp\left(-\left(\frac{(a-A)}{\sigma_c}\right)^2\right)}{K_0 \exp\left(-\left(\frac{(a-a^*)}{\sigma_K}\right)^2\right)}$ | $\sigma_C = 5, K_0 = 5K,$ $a^* = 0, \sigma_K = 1$ |
| Infection rate | $\eta(a) = \eta_0$ constant | multiple values in $[0, 5\ 10^{-2}]$ |
| Viral death rate | $d_v$ constant | $d_v = 5$ day$^{-1}$ |
| Burst size | $V$ constant | $V = 100$ |
| Probability of GTP release | $\gamma$ constant | $\gamma = 0.01$ |
| Transduction probability | $\psi(a, A) = \psi_0 \exp\left(-\left(\frac{(a-A)}{\sigma_\psi}\right)^2\right)$ | multiple values of $\psi_0$ in $[0, 1]$ $\sigma_\psi = 1$ |
| Mutation standard deviation | $h$ constant | $h = 0.01$ |

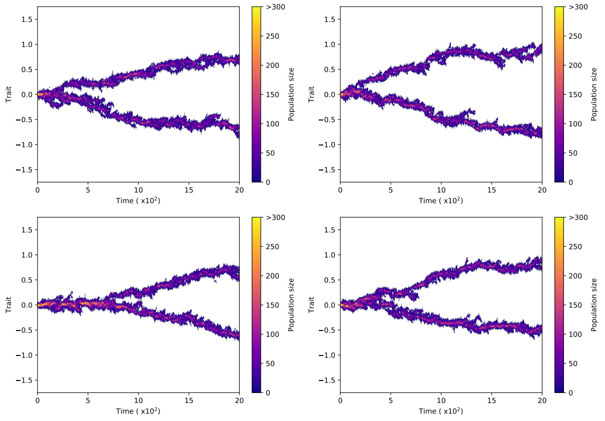

(a) $\psi_0 = 0.$

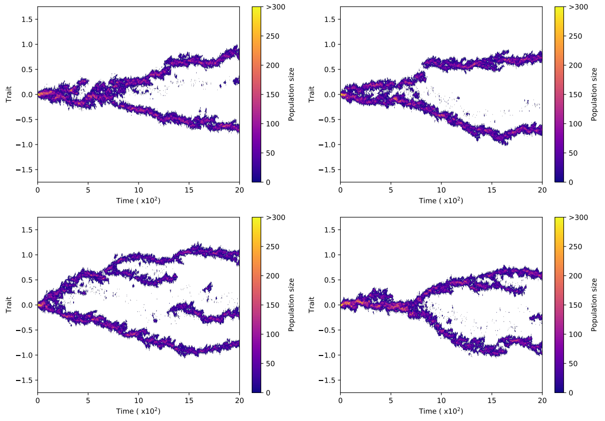

(b) $\psi_0 = 0.01$

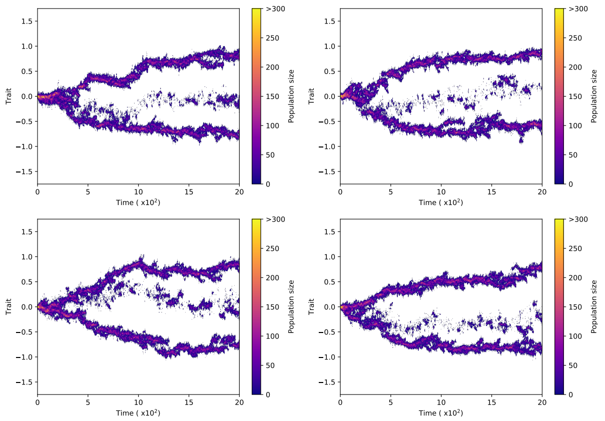

(c) $\psi_0 = 0.05.$

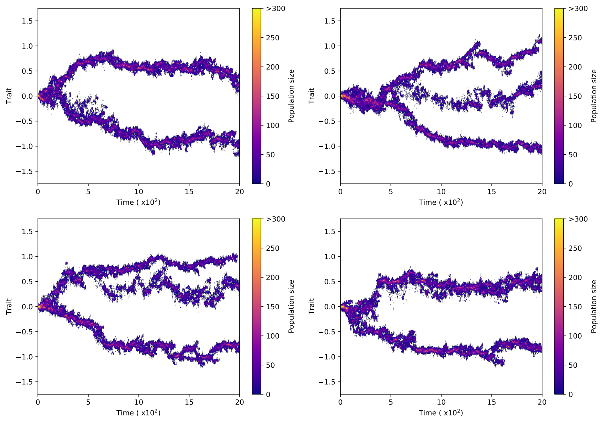

(d) $\psi_0 = 0.1$

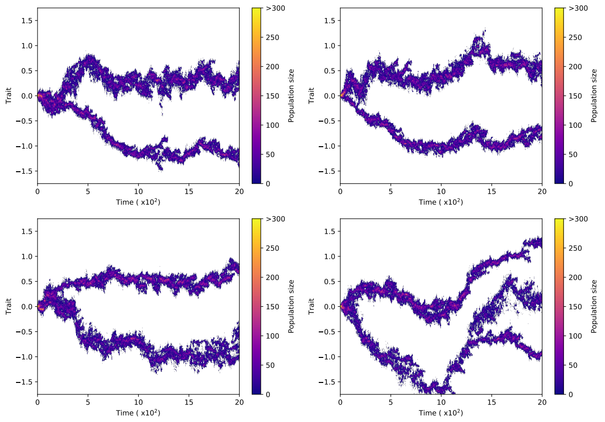

(e) $\psi_0 = 0.25$

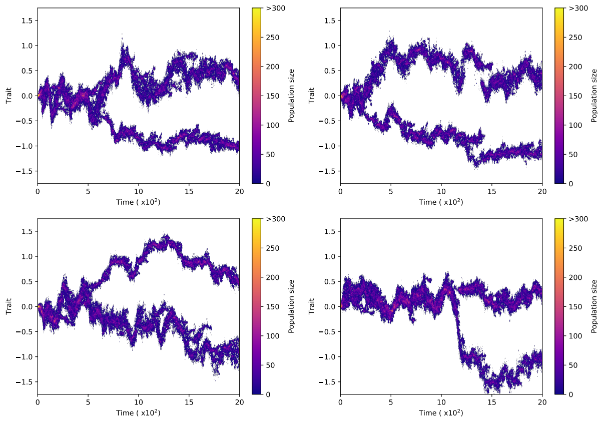

(f) $\psi_0 = 0.5$

Figure 1

**Supplementary figure 1. Individual-based simulations under branching conditions.** Additional examples for each value of $\psi_0$ used in Figure 4 of the main text.

[5] René Descartes. *Discours de la méthode – La Géométrie*. 1637.

[6] Paul Dupuis and Richard S Ellis. *A weak convergence approach to the theory of large deviations*. John Wiley & Sons, 1997.

[7] Jin Feng and Thomas G Kurtz. *Large deviations for stochastic processes*. 131. American Mathematical Soc., 2006.

[8] Mark Iosifovich Freidlin and Alexander D Wentzell. *Random perturbations*. Springer, 1998.

[9] Stefan AH Geritz. "Resident-invader dynamics and the coexistence of similar strategies". In: *Journal of Mathematical Biology* 50.1 (2005), pp. 67–82.

[10] Stefan AH Geritz et al. "Invasion dynamics and attractor inheritance". In: *Journal of Mathematical Biology* 44.6 (2002), pp. 548–560.

[11] Nobuyuki Ikeda and Shinzo Watanabe. *Stochastic differential equations and diffusion processes*. Vol. 24. Elsevier, 2014.

[12] Sylvie Méléard. *Modèles aléatoires en Ecologie et Evolution*. Springer, 2016.

[13] Sylvie Méléard and Vincent Bansaye. *Stochastic models for structured populations*. Springer, 2015.
